# Ribosome-inactivation by a class of widely distributed C-tail anchored membrane proteins

**DOI:** 10.1101/2024.08.16.608256

**Authors:** Robert Njenga, Julian Boele, Friedel Drepper, Kasturica Sinha, Eirini Marouda, Pitter F. Huesgen, Crysten Blaby-Haas, Hans-Georg Koch

## Abstract

Ribosome hibernation is a commonly used strategy that protects ribosomes under unfavorable conditions and regulates developmental processes. Multiple ribosome-hibernation factors have been identified in all domains of life, but due to their structural diversity and the lack of a common inactivation mechanism, it is currently unknown how many different hibernation factors exist. Here, we show that the YqjD/ElaB/YgaM paralogs, initially discovered as membrane-bound ribosome binding proteins in *E. coli,* constitute an abundant class of ribosome-hibernating proteins, which are conserved across all proteobacteria and some other bacterial phyla. Our data demonstrate that they inhibit *in vitro* protein synthesis by interacting with the 50S ribosomal subunit. *In vivo* cross-linking combined with mass spectrometry reveals their specific interactions with proteins surrounding the ribosomal tunnel exit and even their penetration into the ribosomal tunnel. Thus, YqjD/ElaB/YgaM inhibit translation by blocking the ribosomal tunnel and thus mimic the activity of antimicrobial peptides and macrolide antibiotics.

## Introduction

Unicellular organisms, such as bacteria, are constantly challenged by fluctuations in their immediate environment. Consequently, bacteria have developed sophisticated strategies for sensing environmental conditions and for converting this information into metabolic responses ^1–3^. In bacteria, these responses are primarily controlled by transcriptional regulators that adjust gene expression in response to intra- and extracellular cues ^4–7^. These transcriptional control mechanisms are complemented by multiple post-transcriptional response strategies, such as stress-induced adaptation of ribosome biogenesis and protein synthesis ^8–11^. Important players of this adaptation are ribosome-inactivating proteins, which are present in all domains of life and which can inactivate ribosomes either reversibly or irreversibly ^18–22^. Examples are RNA-specific N-glycosidases, such as the plant toxin ricin or the bacterial Shiga toxin, which inactivate ribosomes by depurinating adenine nucleotides within the sarcin-ricin RNA loop of the 23S rRNA in bacteria or the 28S rRNA in eukaryotes ^23,24^. This irreversible ribosome inactivation primarily serves as a defense mechanism against predators or competitors, while reversible ribosome inactivation causing transient ribosome hibernation is used as an important strategy for stress adaptation and for regulating developmental processes ^1,20,25–29^. The *E. coli* ribosome-modulation factor (RMF) and the hibernation-promoting factor (HPF) are well studied bacterial hibernation factors ^30,31^. RMF binds to the 30S ribosomal subunit where it interacts with the ribosomal protein bS1 and induces the dimerization of two 70S ribosomes to an instable 90S dimer, which is subsequently converted into the stable 100S dimer by HPF^32–36^. In stationary *E. coli* cells, approx. 40 to 80% of all ribosomes appear to exist as silent 100S ribosomes ^32,37,38^. Silencing ribosomes protects them against RNAse-dependent degradation ^39^ and adjusts the overall protein synthesis to available nutrients ^19,26,31^. Upon nutrient re-supply, the 100S ribosomes are converted within minutes to active 70S ribosomes ^18,40^. While RMF is only found in γ-proteobacteria, HPF homologues are found in almost all bacteria ^20^. *E. coli* contains additional putative ribosome-inactivating proteins, such as the short HPF-paralog RaiA (ribosome-associated inhibitor A, also referred to as YfiA)^36^, Sra (stationary-phase-induced ribosome associated)^19,41,42^ or RsfS (ribosome silencing factor S)^43^, which are like RMF and HPF soluble proteins that are primarily expressed during stationary phase ^22^.

Furthermore*, E. coli* contains three paralogous membrane-anchored proteins, YqjD, ElaB and YgaM that are shown or predicted to interact with ribosomes during stationary phase. YqjD, ElaB and YgaM belong to the small number of C-tail anchored membrane proteins in *E. coli* ^44–46^. Their production is controlled by the stationary phase specific σ-factor RpoS ^1^ and YqjD was shown to interact with 70S and 100S ribosomes during stationary phase ^45^. However, whether these proteins influence ribosomal activity is not known. Here, we show that YqjD/ElaB/YgaM constitute a widely distributed class of membrane-bound ribosome hibernation factors. YqjD and ElaB inactivate ribosomes by binding to proteins surrounding the ribosomal tunnel exit and even partially protrude into the ribosomal tunnel. Thus, they inactivate ribosomes likely by blocking the ribosomal tunnel, a mechanism that is also observed for macrolide antibiotics and some antimicrobial peptides.

## Results

### YqjD, ElaB and YgaM paralogs are present in many bacterial phyla

YqjD, ElaB and YgaM are largely uncharacterized proteins with a predicted C-terminal transmembrane domain (TM) (Fig. 1A), and they are suggested to be involved in tethering ribosomes to the cytoplasmic membrane ^44,45^. Based on Alphafold2 structural predictions ^47^, they are largely α-helical proteins with a conserved helix-breaking proline residue immediately following the transmembrane domain (Fig. 1B). Our bioinformatic analyses show that the paralogs exhibit significant sequence conservation with 37% sequence identity between YqjD and ElaB and 42% sequence identity between YqjD and YgaM (Fig. 1B). The sequence identity between ElaB and YgaM is 34% and their similarity is 38%.

**Figure 1.**
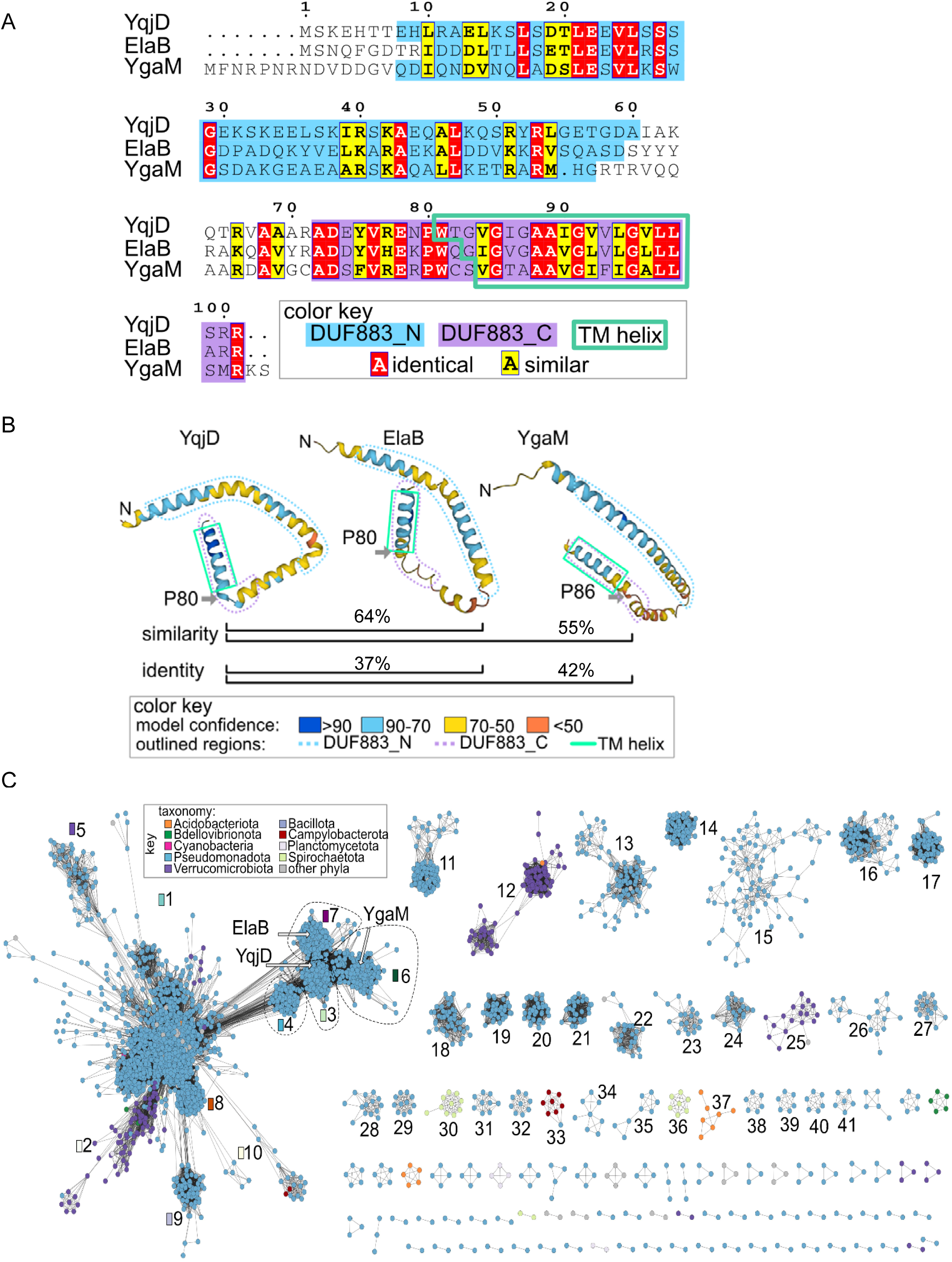
YqjD, ElaB and YgaM represent a widely distributed class of C-tail anchored membrane proteins. (A) Sequence alignment of YqjD, ElaB and YgaM. The locations of DUF883 regions are shaded according to the color key. The predicted C-terminal transmembrane regions (TM) of each protein are indicated by the green box. (B) AlphaFold2-predicted structural models for YqjD, ElaB and YgaM. The cartoons are colored according to model confidence, *i.e.* pLDDT that corresponds to the model’s prediction of its score on the local Distance Difference Test. Lines at the bottom are used to indicate pairwise amino acid similarity and identity. Dotted lines encircle the DUF883 regions (colored according to the key). Green boxes are used to indicate the location of predicted TM regions. (C) Sequence similarity network representing proteins containing DUF883. Nodes are colored by phylum according to the color key. Clusters of proteins containing at least six nodes are labeled. Dotted lines are used to delineate the clusters labelled “SSN cluster” in Fig. 2B. Node information can be found in Supplemental data file S1.

YqjD, ElaB, and YgaM contain DUF883 domains ^48^ (IPR043604: DUF883, N-terminal domain, IPR043605: DUF883, C-terminal domain, and the overlapping IPR010279: Inner membrane protein YqjD/ElaB domain as defined in the InterPro database) that are widespread in different prokaryotic phyla (Figs. 1A &1C, Suppl. Fig. S1). Using a sequence similarity network analysis of proteins matching IPR010279 (which encompasses the N- and C-terminal DUF883 regions), we identified DUF883-containing proteins mainly in Pseudomonadota (9281 proteins), with some homologs found in the PVC group (Planctomycetota-Verruromicrobiota-Chlamydiota (300)), Spirochaetota (130), the delta/epsilon subdivisions (26), Acidobacteriota (19), Campylobacterota (18), Bdellovibrionota (12), and Thermodesulfobacteriota (11) (Fig. 1C). Although some eukaryotic DUF883-containing proteins are predicted, the absence of DUF883-like proteins among closely related eukaryotes suggests that these few eukaryotic proteins could possibly be the result of bacterial contamination in the genome assemblies. In contrast, three separate species from the Archaeal genus *Methanocalculus* contain DUF883 proteins, suggesting that these rare archaeal DUF883 proteins are more likely to be genuine.

Across the YqjD/ElaB/YgaM family, the helix-breaking proline and several glycine residues within the TM helix are highly conserved (Fig. 2A). Analysis of the YqjD, ElaB and YgaM subfamilies reveals the presence of a conserved tryptophan residue that follows the conserved proline, and C-terminal double arginine motifs for YqjD and ElaB, or an arginine-lysine motif for YgaM (Fig. 2A). These putative interfacial residues may stabilize a particular transmembrane orientation ^49^. The TM region is predicted to be slightly longer for YqjD, followed by ElaB, and then YgaM that has the shortest predicted TM helix but the longest C-terminal periplasmic tail (Fig. 1A). Patches of conserved residues among each YqjD/ElaB/YgaM subfamily were also identified (Fig. 2A) as well as residues that have been maintained since the last common ancestor (Suppl. Fig. S2 & S3).

**Figure 2.**
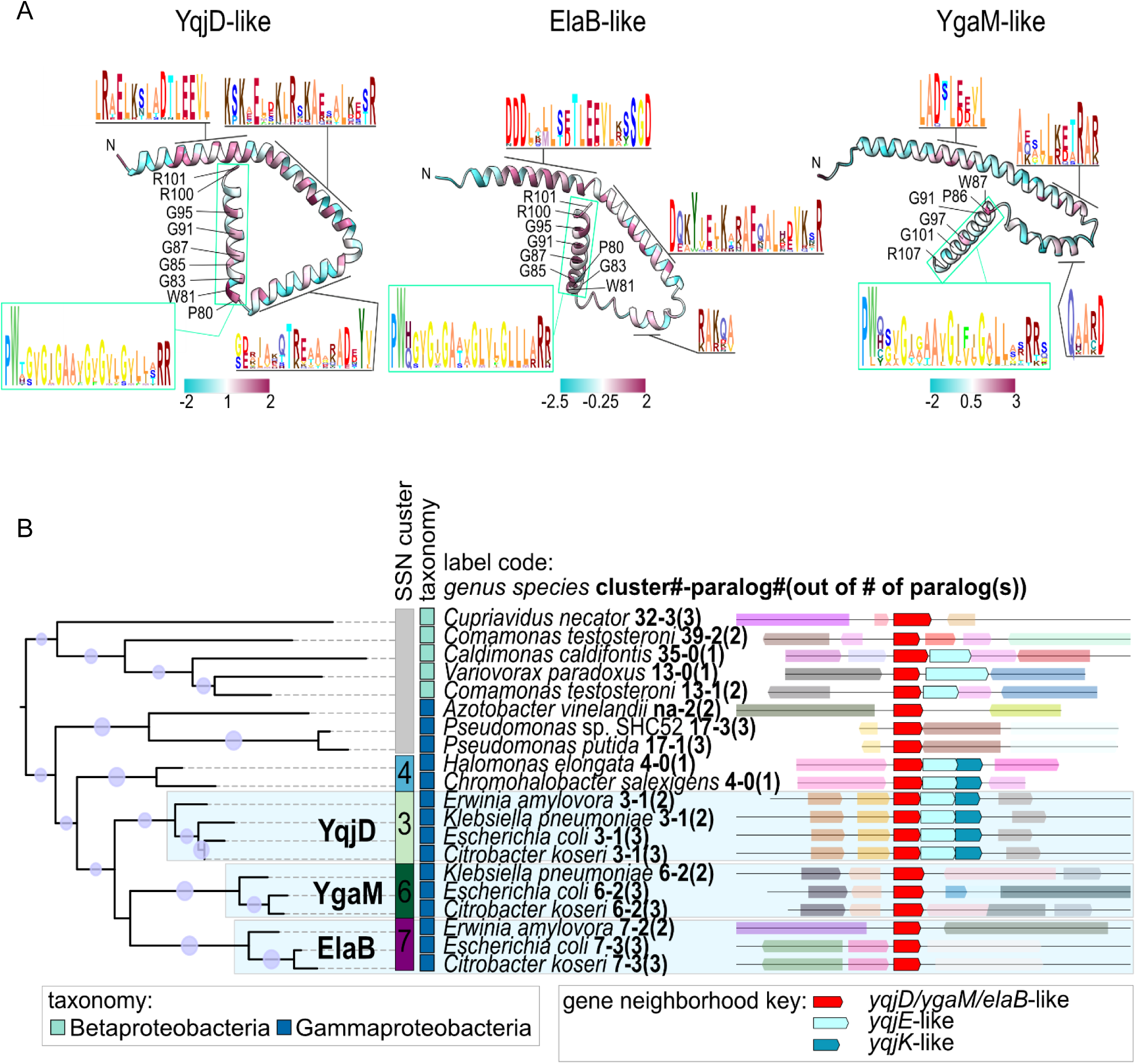
YqjD, ElaB and YgaM represent three distinct subfamilies conserved in Enterobacterales. (A) AlphaFold2-predicted models of YqjD, ElaB and YgaM colored according to sequence conservation in the multiple sequence alignment of proteins from the YqjD, ElaB and YgaM subfamilies. Regions of higher conservation, as well as the transmembrane (TM) region (green box), are shown as sequence logos. Colors represent the conservation values according to the keys. (B) Phylogenetic reconstruction of representative DUF883-containing proteins in beta- and gamma-proteobacteria under maximum likelihood. The “SSN cluster” column corresponds to clusters 3, 4, 6 and 7 in Fig. 1. A complete phylogenetic reconstruction of all clusters is shown in Suppl. Fig. S3A. Taxonomic classification for each node is given according to the color key. The protein labels for each leaf list the organism, followed by the cluster number corresponding to Fig. 1C, followed by the paralog number and followed by the total number of identified paralogs for that organism. If the paralog number is “0’, then that organism has a single DUF883-containing protein, where as “1” indicates 1 of N paralogs. Bootstrap values greater than 0.5 are represented with a normalized purple circle according to the key. The YqjD, ElaB and YgaM clades are shaded with a light blue. For each protein in the gene neighborhood is given. Genes are colored according to shared domains; light pink genes encode proteins with no identified domain. All genes are shown as transparent, except for those genes and homologs listed in the key.

In addition to the taxonomic diversity, our data show that the YqjD/ElaB/YgaM family can be divided into multiple distinct similarity clusters (Fig. 1C, Suppl. Fig. S3). At the similarity threshold used to build the network, YqjD, ElaB and YgaM are found in each of three distinct regions of the network connected to a large cluster dominated by sequences from Pseudomonadota, with numerous smaller protein clusters representing more divergent sequences (Fig. 1C). Based on phylogenetic reconstruction of representatives from each cluster, the family is relatively complex with multiple possible instances of gene duplication, gene loss, and horizontal gene transfer (Fig. 2B, Suppl. Fig. 2 & 3). Based on the conserved presence of *yqjK*-like and/or *yqjE*-like genes, next to genes encoding DUF883 throughout the family (*i.e.*, forming syntenic blocks) (Fig. 2B, Suppl. Fig. S3), YqjD likely represents the archetype of the DUF883 family. The *yqjK* and *yqjE* genes encode for predicted membrane proteins of unknown function. The ElaB and YgaM subfamilies appear to have evolved by subsequent duplication events in Enterobacterales (Suppl. Fig. S2). One early duplication resulted in the YqjD clade and the common ancestor of YgaM/ElaB. A second subsequent duplication event resulted in separate YgaM and ElaB clades (Suppl. Fig. S2). The relatively recent duplication events are evident at the amino acid level, which show significant sequence conservation (Figs. 1B & 2A).

### Growth-phase dependent expression of the paralogous proteins YqjD, ElaB and YgaM

Synthesis of YqjD, ElaB and YgaM is regulated by the σ-factor RpoS and the transcriptional regulator CsrA ^45,50^, indicating that they are produced during stationary phase ^45^. In case of YqjD and ElaB, this is supported by mass spectrometry data, which show increased abundance during stationary phase (Supp. Fig. S3C) ^22,51^. In contrast, the abundance of YgaM was generally lower and MS-data did not reveal increased steady-state levels of YgaM in stationary phase (Supp. Fig. S3C) ^51^. For validating the growth-phase dependent protein levels of YqjD, ElaB and YgaM, we generated peptide antibodies for western blotting with the corresponding knock-out strains as controls. YqjD was detected in wild-type cells after 4h of growth (corresponding to approx. OD_600_ = 3.0) and its levels stayed constant up to 12h (OD_600_ ∼ 5.0) (Fig. 3A). ElaB was also detected after 4h and remained largely unchanged up to 12h (Fig. 3B). The endogenous YgaM levels could not be reliably detected with the available peptide antibodies, which probably reflects its low copy number (Suppl. Fig. S3C) ^51^. In summary, synthesis of YqjD and ElaB starts at late exponential/early stationary phase and their steady-state levels remain stable for several hours.

**Figure 3.**
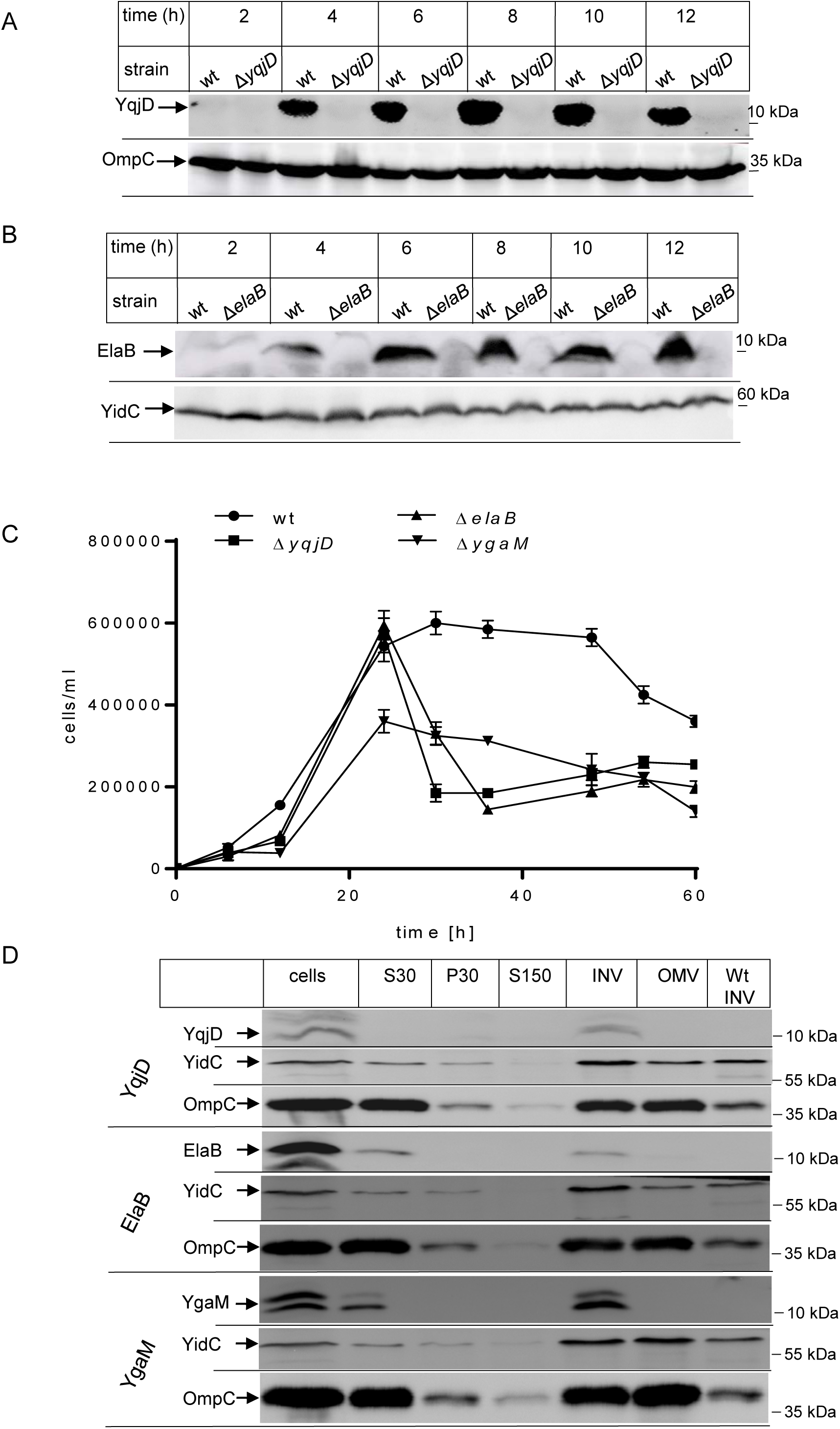
YqjD, ElaB, and YgaM are inner membrane proteins and produced in late exponential phase. (A) *E. coli* wild-type cells BW25113 and the corresponding Δ*yqjD* strain were grown on LB-medium and at the indicated times, 1 x 10^8^ cells were precipitated by 5% trichloroacetic acid (TCA) and loaded on a 15 % SDS-PAGE. The membrane after western blotting was decorated with peptide antibodies against YqjD and OmpC as loading control. (B) As in (A), but wild type and Δ*elaB* cells were analyzed with peptide antibodies raised against ElaB. α-YidC antibodies served as a control. (C) Long-term growth experiment of wt, Δ*yqjD*, Δ*elaB*, and Δ*ygaM* cells on LB medium. Cell counting was performed by the QUANTOM Tx^TM^ Microbial Cell Counter using the QUANTOM viable staining kit, which is an image-based automatic cell counting system that detects individual viable bacterial cells. Indicated are viable cells/ml over time. (D) *E. coli* BW25113 cells expressing His-tagged versions of YqjD, ElaB or YgaM in plasmid pBAD24 were grown to exponential phase and subsequently fractionated after cell breakage. Aliquots of the different fractions were then separated by SDS-PAGE and decorated after western blotting with α-His antibodies. Antibodies against the inner membrane protein YidC and the outer membrane protein OmpC served as controls. S30 and P30 correspond to the supernatant and pellet after a 30.000 x g centrifugation step following cell breakage. The S30 supernatant was then further separated via a 150.000 x g centrifugation into the S150 supernatant and the P150 pellet, the latter was further separated via sucrose gradient centrifugation into the inner membrane vesicle fraction (INV) and the outer membrane fraction (OMV). INVs of the wild type *E. coli* BW25113 served as control. In YgaM-producing cells, α-His antibodies recognized a double band, but it was not further analyzed whether this reflects partial proteolysis.

For monitoring the importance of YqjD/ElaB/YgaM for growth and survival during stationary phase, we performed a long-term growth experiment by counting the viable cell number over 60h on LB-medium for wt and the *yqjD/elaB/ygaM*-deletion strains. The initial growth rate of all four strains was largely comparable, with the exception of the Δ*ygaM* strain, which showed a reduced growth in comparison to the wt (Fig. 3C). Importantly, while the cell number of wild type cells stayed almost constant up to 48h of growth before the cell number declined, the three deletion strains showed a rapid decline already after 20h (Fig. 3C). This supports a particular function of YqjD/ElaB/YgaM during the transition into and during the stationary phase.

Although the generated peptide antibodies recognized YqjD and ElaB in *E. coli* cell extracts, the detection required either a very long exposure (YqjD) or the antibodies recognized additional low-molecular weight bands (ElaB) (Figs. 3A & B). In addition, YgaM could not be reliably detected in whole cell extracts. We therefore constructed copies of *yqjD*, *elaB* and *ygaM* with an N-terminal Xpress-His_6_-tag in the arabinose-inducible pBAD24 vector ^52^. Wild-type cells expressing these constructs were then grown in the presence of arabinose and fractionated by differential centrifugation. All three proteins were recognized by α-His antibodies exclusively in the inner membrane vesicle (INV) fraction, but not in the outer membrane fraction (OMV). As controls, α-YidC and α-OmpC antibodies detected the localization of the inner membrane protein YidC and the outer membrane protein OmpC, respectively. As an additional control, INV of wild-type cells without plasmid were analyzed. This confirms that YqjD, ElaB and YgaM are inner membrane proteins (Fig. 3D).

### YqjD, ElaB and YgaM tether the large ribosomal subunit to the membrane

Whether the YqjD levels influenced the amount of ribosomes bound to INVs was probed with antibodies against ribosomal proteins. Antibodies against the 30S ribosomal protein uS5 did not reveal large differences in the amount of uS5 bound to wild type INVs, INVs from the *yqjD*-expressing strain or the Δ*yqjD* strain (Fig. 4A). In contrast, INVs from the *yqjD-* expressing strain contained increased levels of the 50S ribosomal protein uL2 in comparison to wild-type INVs or Δ*yqjD* INVs. Sucrose gradient purified wild-type ribosomes, containing 70S, 50S and 30S particles, and antibodies against the inner membrane protein YidC served as controls.

**Figure 4.**
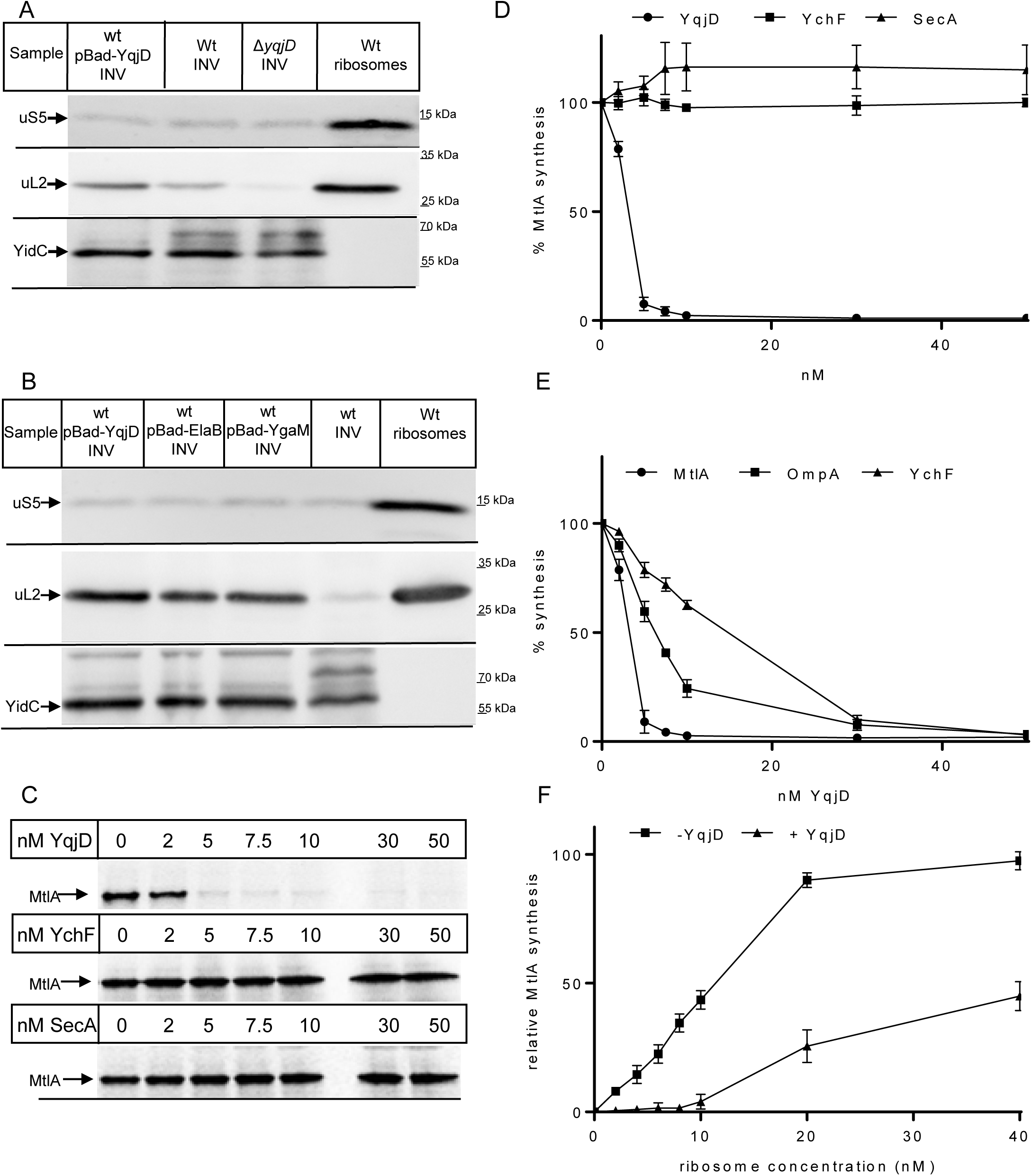
YqjD, ElaB and YgaM are ribosome-inactivating proteins. (A) The inner membrane fractions (INV) of wild type cells (wt), wild type cells containing pBAD-YqjD and Δ*yqjD* cells were separated on SDS-PAGE and after western blotting decorated with antibodies against the 30S ribosomal protein uS5 or the 50S ribosomal protein uL2. Antibodies against the inner membrane protein YidC and sucrose-gradient purified wild type ribosomes served as a control. (B) The INV fractions of wt cells and wt cells containing either pBAD-YqjD, pBAD-ElaB or pBAD-YgaM were processed and controlled as in (A). (C) YqjD was solubilized from *E. coli* membranes, purified via affinity chromatography and added to a cell-free *E. coli in vitro* transcription/translation system containing 30 nM ribosomes ^55^. *In vitro* synthesis of the model protein mannitol permease (MtlA) was performed in the presence of ^35^S-labelled cysteine and methionine. MtlA synthesis in the presence of detergent-containing buffer served as reference (0 nM YqjD). As further controls, the effect of the purified ribosome-interacting proteins YchF and SecA on MtlA synthesis in the *in vitro* system was analyzed. For these controls, detergent-free buffer was used as reference. Samples were separated by SDS-PAGE and analyzed by phosphorimaging. (D) Quantification of four independent experiments as shown in (C). The amount of MtlA synthesized was quantified using a phosphorimager and the *Image1/Fiji* software and synthesis in the absence of YqjD, YchF, or SecA, respectively, was set to 100%. The values on the X-axis refer to the final concentration (nM) of the added protein (YqjD, YchF, or SecA) in the cell-free transcription/translation system. (E) As in (C) and (D), but *in vitro* synthesis of the outer membrane protein OmpA and the cytosolic protein YchF were also tested in the presence of increasing YqjD concentrations. (F) *In vitro* protein synthesis of MtlA was performed either in the absence of YqjD (-YqjD) or in the presence of 30 nM YqjD (+ YqjD) and increasing concentrations of sucrose-gradient purified ribosomes. Quantification after SDS-PAGE and phosphorimaging was performed as in (D) and (E) and MtlA synthesis at 40 nM ribosomes was set to 100%. The error bars indicate the SD.

We also tested the localization of the 30S and 50S subunits in INVs of the *elaB* and *ygaM* overexpressing strains. As seen for YqjD, the amounts of uS5 in these INVs were comparable to wild-type INVs, but the levels of uL2 were increased (Fig. 4B). These data indicate that YqjD, ElaB and YgaM preferentially tether the 50S ribosomal subunit to the membrane, but not the 30S subunit. However, this does not exclude that YqjD/ElaB/YgaM initially bind to 70S ribosomes, which then dissociate during INV preparation.

The interaction between YqjD and ribosomes was further studied in an *in vitro* approach using purified YqjD and sucrose gradient-purified ribosomes isolated from cells grown to either exponential or stationary phase. This revealed binding of YqjD to both types of ribosomes, although binding to stationary phase ribosomes was slightly less efficient (Suppl. Fig. S4A). *E. coli* ribosomes isolated via sucrose-gradient centrifugation are almost exclusively non-translating ^11,53^, showing that YqjD primarily interacts with non-translating ribosomes.

If YqjD inactivates ribosomes, high concentrations of YqjD should reduce the number of translation-competent ribosomes and increase chloramphenicol sensitivity, because chloramphenicol targets translating ribosomes by inhibiting their peptidyltransferase activity. The minimal inhibitory concentration of chloramphenicol is in the range of 20-30 µg/ml ^54^, however, *yqjD*-expressing cells were already inhibited at 0.75 µg/ml (Suppl. Fig. S4B). This hypersensitivity of *yqjD-*expressing cells against chloramphenicol supports a possible role of YqjD in ribosome hibernation. We did not observe increased chloramphenicol resistance in the Δ*yqjD* strain, which is likely explained by the surplus of ribosomes over YqjD in *E. coli* cells.

### YqjD prevents translation *in vitro*

For validating whether YqjD interferes with translation, we employed a coupled *in vitro* transcription/translation system, consisting of a purified cell extract (CTF, cytosolic translation factors) and purified ribosomes ^55^. Adding increasing amounts of purified YqjD to the *in vitro* system inhibited synthesis of mannitol permease (MtlA), which is frequently used as model membrane protein for *in vitro* studies ^56^ (Fig. 4C). This was different for two additional ribosome-interacting proteins, YchF and SecA^11,57,58^, which did not inhibit MtlA synthesis (Figs. 4C & 4D). YqjD also inhibited *in vitro* synthesis of the cytosolic protein YchF and the secretory protein OmpA (Fig. 4E), although the inhibitory effect was slightly less pronounced than the inhibition of MtlA synthesis. This could relate to differences in the *in vitro* translation speed due to codon usage or mRNA length ^59^. Finally, a concentration dependent inhibition of MtlA synthesis was also observed with ElaB, although full inhibition required higher concentrations than in the case of YqjD (Supp. Figs. S5A & B).

If YqjD stoichiometrically inactivates *E. coli* ribosomes, this effect should depend on the ribosome concentration in the *in vitro* assay. The *in vitro* system routinely contains 10-30 nM ribosomes ^55^, although there are variations in the translational activities between different ribosome preparations. MtlA synthesis was analyzed at different ribosome concentrations in the absence or presence of 30 nM YqjD. In the absence of YqjD, MtlA synthesis increased with increasing ribosome concentrations and reached saturation at approx. 20 nM ribosomes (Fig. 4F). In the presence of YqjD, synthesis was strongly reduced up to approx. 10 nM ribosomes, but then gradually increased (Fig. 4F). At approx. 20 nM ribosomes, MtlA synthesis was detectable even in the presence of YqjD. These data demonstrate that YqjD prevents translation by inactivating ribosomes.

### Ribosome-inactivation by YqjD depends on the N-terminus and on dimerization

Previous data had indicated that YqjD interacts via its N-terminus with ribosomes ^45^ and we therefore tested MtlA synthesis in the presence of N-terminally truncated YqjD variants. *In vitro* MtlA synthesis was almost completely blocked by the addition of 5 nM full-length YqjD (Fig. 5A). In contrast, gradually deleting N-terminal amino acids diminished YqjD’s ability to prevent MtlA synthesis (Fig. 5A), demonstrating that the N-terminus is required for ribosome inactivation. Surprisingly, deleting the C-terminal transmembrane domain (ΔTM-YqjD) also significantly reduced the inhibitory effect of YqjD on translation (Fig. 5A). The *in vitro* translation assays were performed in the presence of purified YqjD, but in the absence of membranes, demonstrating that membrane tethering of ribosomes cannot be responsible for impaired translation in our assay. Adding the purified transmembrane helix of YqjD to the *in vitro* synthesis assay did also not reduce MtlA synthesis (Fig. 5B), indicating that the TM is only indirectly involved in ribosome inactivation. Whether the lack of the TM influenced ribosome binding was analyzed by incubating purified YqjD, ΔTM-YqjD and ΔN(12)-YqjD with purified ribosomes. In comparison to full-length YqjD, both truncated variants showed reduced ribosome binding (Suppl. Fig. S6A), but the truncation of the N-terminus had a stronger effect than the deletion of the C-terminus. Thus, even in the absence of the TM, YqjD retains some ribosome binding activity but completely fails to inactivate ribosomes.

**Figure 5:**
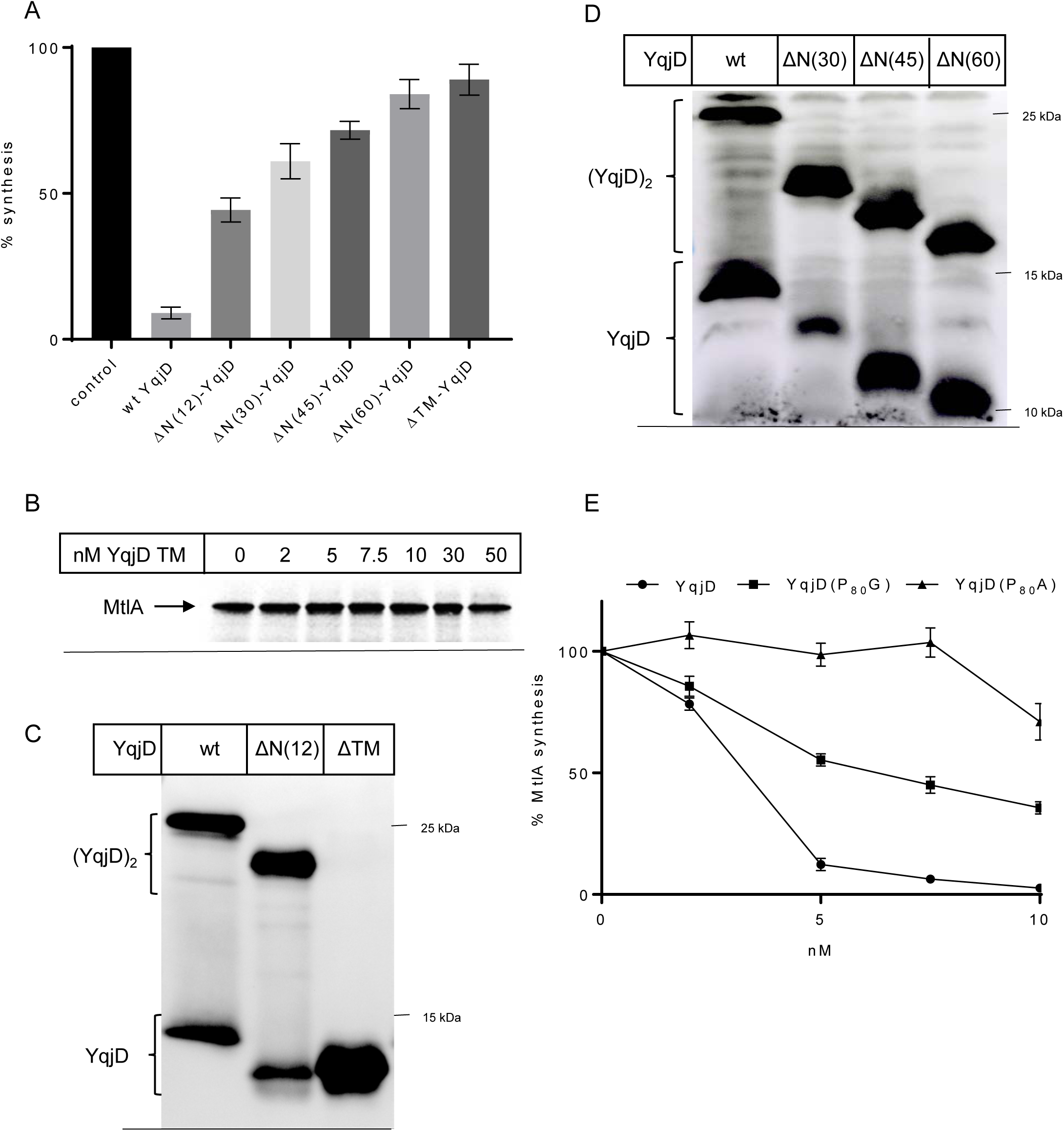
Both N- and C-terminus of YqjD are required for inhibiting protein synthesis. (A) *In vitro* MtlA synthesis was performed as described in the legend to Fig. 4 and analyzed in the absence of YqjD (control) and in the presence of 5 nM wild-type YqjD, or YqjD variants that were N-terminally truncated or lacked the C-terminal 18 amino acids (ΔTM-YqjD). MtlA synthesis in the absence of YqjD was set to 100 %. Shown are the mean values of 3 independent experiments and the error bar reflect the SD. (B) The TM helix of YqjD was chemically synthesized and added in increasing concentrations to the MtlA synthesizing *in vitro* translation system. Samples were separated by SDS-PAGE and analyzed by phosphorimaging as in (A). (C) Purified YqjD, ΔN(12)-YqjD and ΔTM-YqjD were denatured in loading dye at 56 °C, separated on SDS-PAGE and probed with α-Xpress antibodies. (D) As in (C), but ΔN(30), ΔN(45) and ΔN(60) were analyzed and compared to full-length YqjD (wt). (E) Purified YqjD and its variants in which the proline residue at position 80 was replaced by either glycine or alanine were added in different concentrations to the MtlA synthesizing *in vitro* translation system. After SDS-PAGE and phosphorimaging, the MtlA synthesis in the absence of YqjD was set to 100 %. Shown are the quantifications of three independent experiments and the error bars reflect the SD. The values on the X-axis refer to the final concentration (nM) of the added protein (YqjD, YqjD(P_80_G), or YqjD(P_80_A)) in the *in vitro* transcription/translation system.

YqjD contains several glycine residues in its transmembrane domain (Fig. 2A). Such glycine motifs have been shown to promote homo-dimer formation ^60^ and indeed, YqjD shows a strong propensity for dimer formation, which are stable even on SDS-PAGE (Fig. 5C). Dimerization of ElaB and YgaM was less pronounced, but this could simply reflect reduced stability of the dimer on SDS-PAGE (Suppl. Fig. S5C). YqjD dimerization was not influenced by the His-tag, because the tag-free YqjD showed a comparable amount of dimer (Suppl. Fig. S6B). Interestingly, YqjD dimerization was completely diminished when the C-terminal TM was deleted (Fig. 5C), while the N-terminal truncations had no influence on dimerization (Figs. 5C & 5D). Thus, dimerization of YqjD depends on the TM and the lack of ribosome inactivation by ΔTM-YqjD could reflect its failure to dimerize. This was further tested by constructing YqjD variants, in which the native TM was replaced by the N-terminal TM of either FtsQ or YfgM or by the C-terminal TM of the mitochondrial protein Fis1. These variants showed the same inhibitory effect on MtlA synthesis as wild-type YqjD (Suppl. Fig. S6C), and also formed SDS-resistant dimers, although dimer stability on SDS-PAGE was reduced in comparison to the native YqjD (Suppl. Fig. S6D). Thus, the nature of the TM is not important for ribosome inactivation. Instead, the data rather indicate that the TM is required for YqjD dimerization, which in turn promotes ribosome inactivation via the N-terminus.

Dimerization could induce ribosome inactivation by reducing the topological flexibility of the N-terminus, which potentially also explains the conservation of the proline residue in immediate vicinity to the TM of YqjD. Proline residues can act as α-helix breaking amino acids, which promote rigid body motions of helical segments ^61^. This is visible in the Alphafold structural prediction of YqjD (Fig. 1B), which shows that the proline residue induces a strong kink after the TM. The importance of the proline residue was tested by replacing it with either glycine, which has a lower helix-breaking propensity than proline, or with alanine, which stabilizes the α-helix. The YqjD(P_80_G) variant prevented MtlA synthesis to a lesser extent than wild-type YqjD (Fig. 5E), and the inhibitory activity was even further reduced in the YqjD(P_80_A) variant (Fig. 5E). In summary, ribosome inactivation by YqjD is likely dependent on the correct orientation of its N-terminus, which is compromised when the C-terminal TM is deleted or when the conserved proline residue is mutated.

### YqjD contacts ribosomal proteins close to the peptide tunnel

For determining the YqjD-ribosome contacts in more detail, we used an *in vivo* site-directed cross-linking approach. The UV-sensitive phenylalanine derivative para-benzoyl-L-phenylalanine (pBpa) was site specifically inserted at the N-terminal positions 10 and 39 of YqjD, using an amber-suppressor tRNA and a cognate tRNA synthetase ^62,63^. *E. coli* cells expressing either YqjD(L10pBpa) or YqjD(I39pBpa) were then grown on LB-medium in the presence or absence of pBpa. The two YqjD variants were only visible in the presence of pBpa, demonstrating the successful insertion of pBpa into YqjD (Fig. 6A). The two YqjD variants also showed the same propensity for dimerization as the wild-type YqjD (Fig. 6A). When YqjD(L10pBpa) or YqjD(I39pBpa) expressing *E. coli* cells were exposed to UV-light, multiple additional bands were recognized by α-Xpress antibodies in the UV-treated sample, but not in the sample not exposed to UV-light (Fig. 6B). Some additional bands were also visible in UV-treated wild-type cells, lacking pBpa (Fig. 6B). These pBpa-independent cross-linking products are likely the result of UV-induced radical formation of aromatic amino acids that favor non-specific protein-protein and protein-nucleic acid cross-links ^64,65^. The cross-linked material was then further enriched by affinity chromatography and probed with α-Xpress antibodies. This revealed UV-dependent cross-linked bands at approx. 40 kDa and 55 kDa for YqjD(L10pBpa) and YqjD(I39pBpa), which were not visible for wild-type YqjD lacking pBpa (Fig. 6C). For YqjD(L10pBpa) additional weaker bands at approx. 110 kDa were also visible.

**Figure 6:**
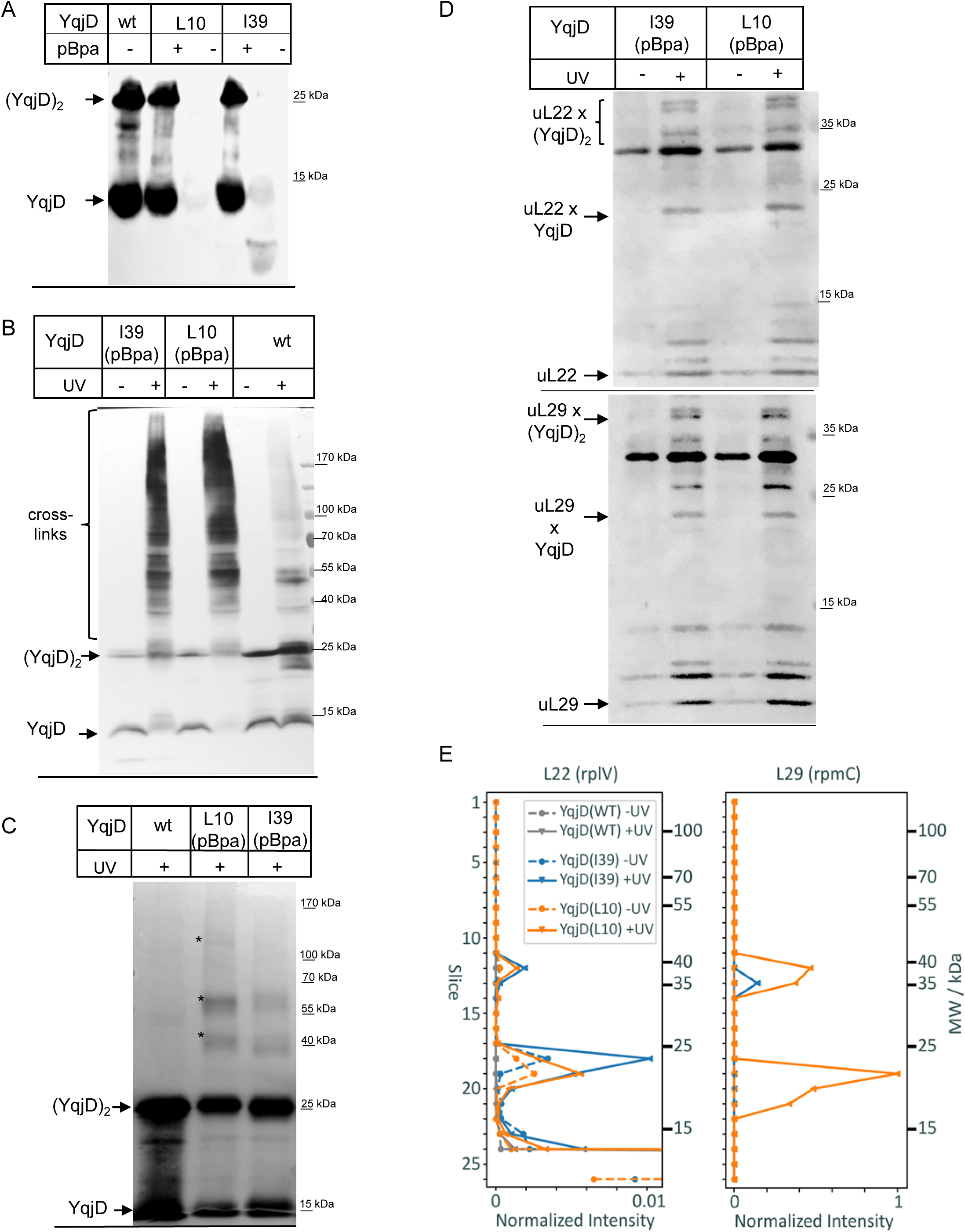
YqjD interacts via its N-terminus with proteins of the ribosomal peptide tunnel. (A) Cells producing either wild-type YqjD or its variants containing the amber stop codons at position 10 or 39 for inserting the UV-sensitive amino acid derivative para-benzoyl-L-phenylalanine (pBpa) were grown in the presence or absence of pBpa. After TCA precipitation of whole cells, the material was separated by SDS-PAGE. After western blotting, the samples were decorated with antibodies against the N-terminal Xpress tag. Indicated are the YqjD monomer and dimer. (B) As in (A), but *E. coli* cells were UV-exposed for inducing the cross-link reaction. After TCA precipitation and SDS-PAGE, the material was decorated with α-Xpress antibodies. (C) As in (B), but after UV exposure of whole cells, YqjD and its cross-linked partner proteins were affinity purified via its His-tag and analyzed after SDS-PAGE by immune detection with Xpress antibodies. (D) The material in (C) was probed with antibodies against the ribosomal protein uL22 (upper panel) or uL29 (lower panel). Indicated are putative cross-links between YqjD and ul22 or uL29, respectively. (E) Affinity-purified YqjD and its cross-linked partner proteins as in (C) were separated on SDS-PAGE and the gel lanes were sliced into multiple slices, which were separately processed and analyzed by mass spectrometry. Shown are the normalized intensities of uL22 and uL29 peptides in the –UV and +UV treated samples of wild-type YqjD, YqjD(I39) and YqjD(L10).

Proteins surrounding the ribosomal tunnel exit on the 50S subunit serve as hot spots for ribosome-interacting proteins ^57,66,67^. As YqjD likely interacts with the 50S ribosomal subunit (Fig. 4A), the cross-link samples were analyzed with antibodies against the ribosomal proteins uL22 and uL29, which are located next to the tunnel exit. uL22 has a MW of 12 kDa and the α-uL22 antibodies recognized several UV-dependent bands; the most prominent migrated just below the 25 kDa marker band and could reflect a cross-link between monomeric YqjD and uL22 (Fig. 6D, upper panel). At approx. 35 kDa, three UV-dependent bands were observed for both YqjD(L10pBpa) and YqjD(I39pBpa), which could reflect a cross-link between dimeric YqjD and uL22 (Fig. 6D, upper panel). The migration of a single cross-linking product in multiple species on SDS-PAGE is often observed ^68–70^ and reflects different three-dimensional shapes of the covalently linked proteins.

The α-uL29 antibodies also recognized several UV-dependent bands, which corresponded in size to crosslinks between the 7.3 kDa uL29 and monomeric and dimeric YqjD. (Fig. 6D, lower panel). Overall, the sequence similarity of ribosomal proteins and the low specificity of the available antibodies did not allow for an unambiguous identification of cross-links between YqjD and 50S ribosomal proteins. The samples after cross-linking were therefore analyzed by mass spectrometry. This identified two UV-dependent uL22 cross-linking products of 23 kDa and 38 kDa, respectively, which were observed for both YqjD(L10pBpa) and YqjD(I39pBpa) (Fig. 6E). This demonstrates that both monomeric and dimeric YqjD is cross-linked to uL22. The MS analyses also revealed two uL29 cross-linking products of YqjD(L10pBpa) at 21 kDa and 37 kDa, while for YqjD(I39pBpa) only one cross-linking product at 35 kDa was detected (Fig. 6E). In addition, the MS revealed single cross-linking products between uL23 and YqjD(I39pBpa) and between uL24 and YqjD(I39pBpa) (Suppl. Fig. 7A), which were not detected by antibodies. Proteins uL23 and uL24 are, like uL29 and uL22, located in close vicinity to the ribosomal tunnel exit and form a platform for multiple ribosome-interacting proteins ^66,67,71,72^. In summary, these data demonstrate that YqjD interacts with 50S ribosomal proteins that surround the ribosomal tunnel exit.

### ElaB prevents translation by mimicking anti-microbial peptides

The contact between YqjD and the ribosomal proteins uL22 and uL23 is intriguing, because both proteins are not only exposed to the ribosomal surface but also contact the interior of the ribosomal tunnel via β-hairpin loops. Subunit uL22 together with uL4 forms a central constriction within the ribosomal tunnel, which serves as a binding site for macrolide antibiotics and antimicrobial peptides ^73–75^. The intra-tunnel loop of uL23 is located closer to the tunnel exit and acts as a nascent chain sensor that binds to the protein targeting factors SRP and SecA ^57,66,67^. Thus, it appeared possible that YqjD interferes with translation by inserting into the ribosomal peptide tunnel and as such mimics antimicrobial peptides ^23,74,76,77^, or eukaryotic ribosome hibernation factors ^29,78^.

For monitoring whether YqjD reaches into the peptide tunnel of the ribosome, ribosomes that contained the cross-linker pBpa at the tip of the intra-tunnel β-hairpin loop of uL23 (position 71) or at position 52 of the surface-exposed globular domain of uL23 were generated, isolated ^57,66^ and incubated *in vitro* with purified YqjD, followed by UV-exposure. As a control, these ribosomes were incubated with purified SRP, which was previously shown to contact both uL23 residues ^66^. While UV-dependent cross-links to Ffh, the protein component of the *E. coli* SRP ^79^, were observed from both residue 71 and residue 52 (Fig. 7A, left panel), no cross-links between uL23 to YqjD were visible in this *in vitro* approach (Fig. 7A, right panel).

**Figure 7:**
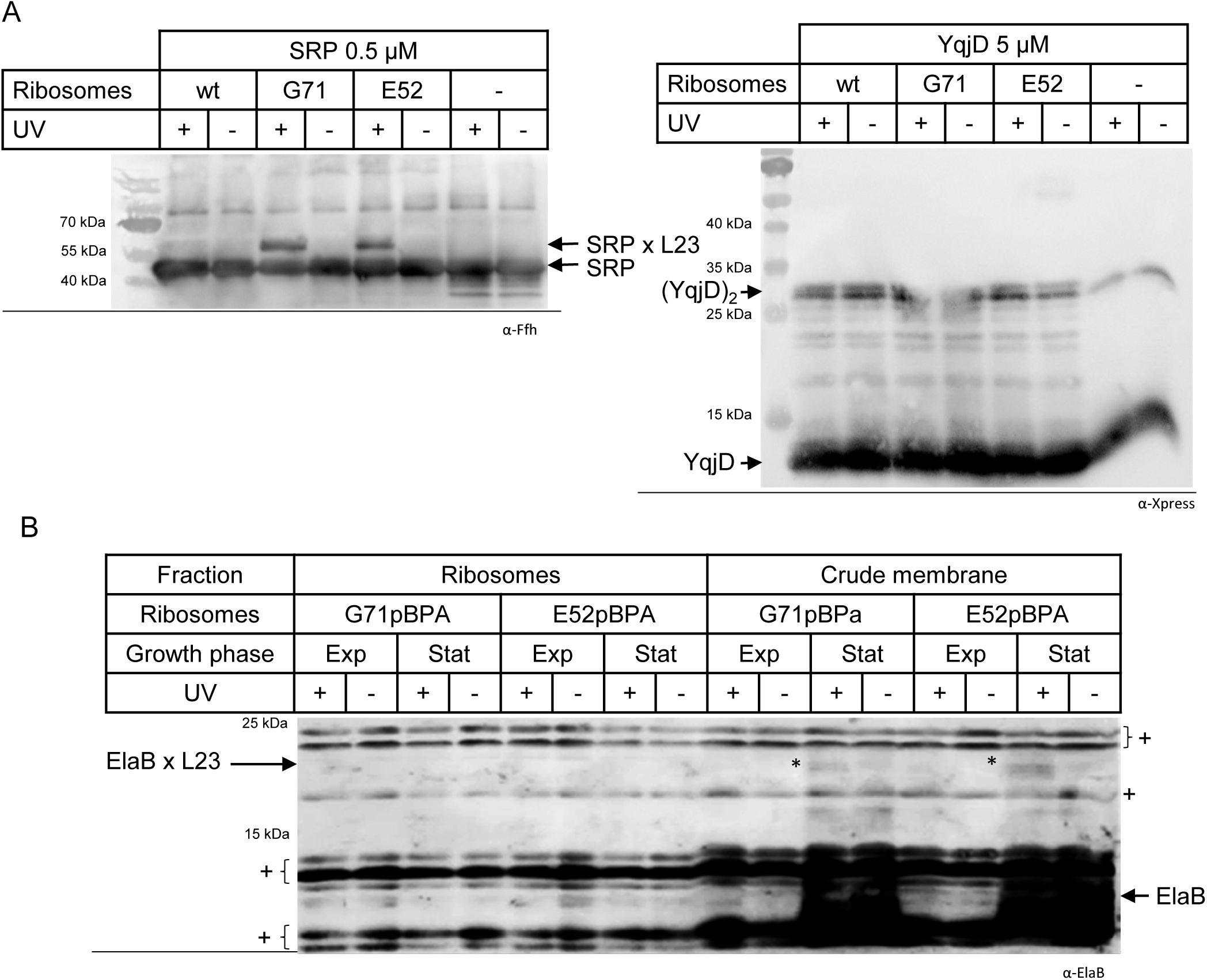
ElaB enters the ribosomal peptide tunnel *in vivo*. (A) *In vitro* site directed cross-linking was performed with ribosomes containing the cross-linker pBpa either at the surface-exposed position 52 of uL23 or at position 71 within the ribosomal peptide tunnel. Ribosomes were purified via sucrose-gradient centrifugation and incubated for 20 min at 30 °C with equimolar amounts (500 nM) of purified signal recognition particle (SRP) or a 10-fold excess of purified YqjD. After incubation, the sample was exposed to UV light and then separated by SDS-PAGE and decorated with antibodies against Ffh, the protein component of the *E. coli* SRP (A, left panel) or antibodies against the Xpress tag of YqjD (A, right panel). Wild-type ribosomes (no pBpa) and samples without ribosomes served as controls. Indicated are the cross-links to SRP from the surface exposed uL23 residue E52 and the intra-tunnel residue G71. (B) *In vivo* site-directed cross-linking was performed with *E. coli* cells producing pBpa-containing ribosomes. The *E. coli* Δ*rplW* (uL23) deletion strain expressing uL23(E52pBpa) or uL23(E71pBpa) was grown to exponential or stationary phase and UV-exposed. After UV exposure of whole cells, cells were fractionated into the soluble ribosome fraction and the crude membrane fraction. These fractions were then separated by SDS-PAGE and probed with peptide antibodies against the native ElaB. The potential cross-links between ElaB and uL23(G71pBpa) and uL23(E52pBpa) are labelled by (*). Note, that the antibody non-specifically recognizes multiple bands, which are indicated by (+).

We therefore switched to an *in vivo* approach. *E. coli* Δ*rplW* cells, which lack the uL23-encoding *rplW* gene on the chromosome but contain either the plasmid-encoded uL23(E52pBpa) or the uL23(G71pBpa) variant, were grown to exponential or stationary phase and then UV-exposed. The *rplW* (uL23) gene is essential in *E. coli,* and cells containing these plasmid-encoded versions were only able to grow in the presence of pBpa, demonstrating that these pBpa-containing ribosomes are functional, which is in agreement with previous reports ^57,66^. After cell lysis, cells were fractionated into the soluble ribosome fraction and the crude membrane fraction, which were then decorated with α-YqjD antibodies. However, the low specificity of these antibodies did not allow us to detect specific cross-linking bands (Suppl. Fig. 7B). In contrast, when the same material was decorated with α-ElaB antibodies, we found weak UV-dependent cross-links at approx. 20 kDa for both position 52 and 71 of uL23 (Fig. 7B, indicated by *), fitting to the size of a ElaB-uL23 cross-link. Importantly, these cross-links were only visible in cells grown to stationary phase, but not exponentially grown cells. This observation is explained by the increased production of native ElaB when cells enter stationary phase (Fig. 3B). As ElaB is a membrane protein (Fig. 3D), the cross-linked band was detected only in the membrane fraction, but not in the soluble ribosome fraction. The cross-linking product is rather weak, but this experiment was executed in the presence of native ElaB, which is greatly sub-stoichiometric to ribosomes *in vivo*. Nevertheless, due to the non-specific recognition of many proteins by the α-ElaB peptide antibody (Fig. 7B, indicated by ^+^) and the lack of detectable YqjD-uL23 cross-links, additional studies need to further validate that ElaB and potentially also YqjD/YgaM inactivate ribosomes by penetrating into the ribosomal peptide tunnel. This would then indicate that YqjD/YgaM/ElaB mimic the strategy of antimicrobial peptides ^74^ and eukaryotic ribosome-hibernation factors, such as Dap1b ^29^ and Mdf2 ^78^. However, considering that the YqjD dimerization appears to be important for ribosome inactivation, the N-termini of a YqjD dimer will not be able to reach deeply enough into the tunnel to reach the peptidyltransferase center of the ribosome. Still, the proximal vestibule of the ribosomal tunnel is likely wide enough to accommodate two N-termini ^80,81^.

## Discussion

Ribosome biogenesis and protein synthesis are the most energy consuming cellular processes and they are therefore strictly regulated in response to nutrient availability and stress conditions ^9,13,22,82,83^. This saves energy and reduces the overall production of damage-prone proteins, while a basal level of protein synthesis is maintained. Complementary to these strategies are ribosome-inactivating mechanisms, which provide the fast and reversible means for shutting down the activity of already assembled ribosomes ^20,21,27,30,41,84–86^. Due to the high structural diversity of hibernation factors and the lack of common mechanisms by which they interfere with ribosomal activity ^19,31^, it is currently unknown how many different hibernation factors exist in bacteria. Recently, a novel hibernation factor, called Balon, was identified in the cold-adapted bacterium *Psychrobacter urativorans* and shown to occupy the ribosomal A-site in complex with EF-Tu ^88^. Balon homologues were found in 23 out of 27 bacterial phyla, but are absent in *E. coli* ^88^.

Previous studies had identified YqjD as an *E. coli* ribosome-interacting protein, which inhibited cell growth when over-produced ^45^. It was therefore suggested that YqjD and its paralogs ElaB and YgaM might act as membrane-bound ribosome-hibernation factors in *E. coli* ^1,18,50^. This was experimentally verified in our study, which demonstrates that YqjD and ElaB inhibit *in vitro* protein synthesis in a dose-dependent manner. Our study also identified YqjD/ElaB/YgaM proteins in many bacterial species, suggesting that they potentially represent a widely distributed family of ribosome hibernation factors. Although sequence similarity can be a poor predictor of biological function, sequence similarity combined with gene neighbor conservation (*yqjE*- and *yqjK*-like genes) suggests that the function of these proteins could be conserved across and outside of the Pseudomonadota phylum. Since there have clearly been multiple instances of gain/loss of paralogs and, potentially, horizontal gene transfer, functional tailoring has occurred in different lineages and species. However, whether such tailoring has affected molecular and/or biological functions of any individual uncharacterized protein is as-of-yet unknown.

Based on sucrose-gradient centrifugation, YqjD was suggested to bind to the 30S ribosomal subunit ^45^. However, our data indicate that YqjD/ElaB/YgaM preferentially interact with the 50S ribosomal subunits and tether them to the *E. coli* membrane. Furthermore, by *in vivo* cross-linking combined with mass spectrometry, we demonstrate that the N-terminus of YqjD interacts with the ribosomal proteins uL22, uL23, uL24 and uL29, which encircle the ribosomal peptide tunnel exit on the 50S ribosomal subunit. Thus, it is possible that YqjD can bind to both ribosomal subunits.

Previous studies have revealed the importance of the ribosomal peptide tunnel as binding site for chaperones and targeting factors. ^57,66,67,89–91^. Eukaryotic dormancy factors ^29,78^ or antimicrobial peptides and macrolide antibiotics ^74,92^ also target the ribosomal peptide tunnel. Intriguingly, the YqjD paralog ElaB, inserts into the ribosomal tunnel and contacts the β-hairpin loop of uL23, which was previously identified as an intra-tunnel nascent chain sensor ^57,66^. The uL23 β-hairpin loop is located in the lower section of the ribosomal tunnel, which is generally wide enough to allow proteins to enter ^91,93^. This has been demonstrated for SRP ^66,94^, SecA ^57^ and the cytosolic loops of SecY ^95,96^. On the other hand, antimicrobial peptides, such as oncocin or bactenecin, have been shown to contact the A-site tRNA binding pocket and the A-site crevice, demonstrating that they deeply insert into the ribosomal tunnel ^97,98^. Whether ElaB is also able to contact the upper section of the tunnel is currently unknown. While ElaB can enter the ribosomal tunnel, we did not observe this for YqjD. Thus, we can currently not exclude that ElaB and YqjD inhibit protein synthesis by different mechanisms, as also deduced from the low sequence conservation of their respective N-termini, differences in dimer stability and lower ribosome inactivation potential of ElaB in comparison to YqjD.

The importance of YqjD’s N-terminus for ribosome binding was already shown previously ^45^ and we demonstrate here that the N-terminal truncated YqjD is impaired in ribosome inactivation. Surprisingly, deleting the C-terminal TM of YqjD also significantly reduced ribosome inactivation. The transmembrane domain of YqjD contains several conserved glycine residues, which are often involved in dimerization or oligomerization ^60,99^. In support of this, our data show that the propensity of YqjD to form stable dimers even on SDS-PAGE is strictly dependent on the TM. The lack of ribosome inactivation when the dimerization-promoting TM of YqjD is missing could indicate that the YqjD-ribosome interaction is primarily avidity-driven. Thus, each YqjD has only a low affinity for ribosomes, but high affinity ribosome binding is achieved when two or more YqjD monomers oligomerizes. Avidity-driven interactions with the ribosome are not unusual and have been shown for example for trigger factor dimers ^100^. This probably also explains the importance of the conserved proline residue. Proline-kinked α-helices have been suggested to form cage- or funnel-like structures ^101^, and their reduced topological flexibility could help to orient multiple N-termini in close proximity to the ribosome. The reduced ribosome binding of the ΔTM-YqjD variant, supports such an avidity-driven interactions. Still, it is surprising that although ΔTM-YqjD retains some residual ribosome binding activity, it completely fails to inactivate ribosomes. Thus, it is possible that ribosome inactivation depends on the simultaneous binding of both N-termini of the YqjD dimer to a single ribosome, but this needs to be further explored (Fig. 8).

**Figure 8:**
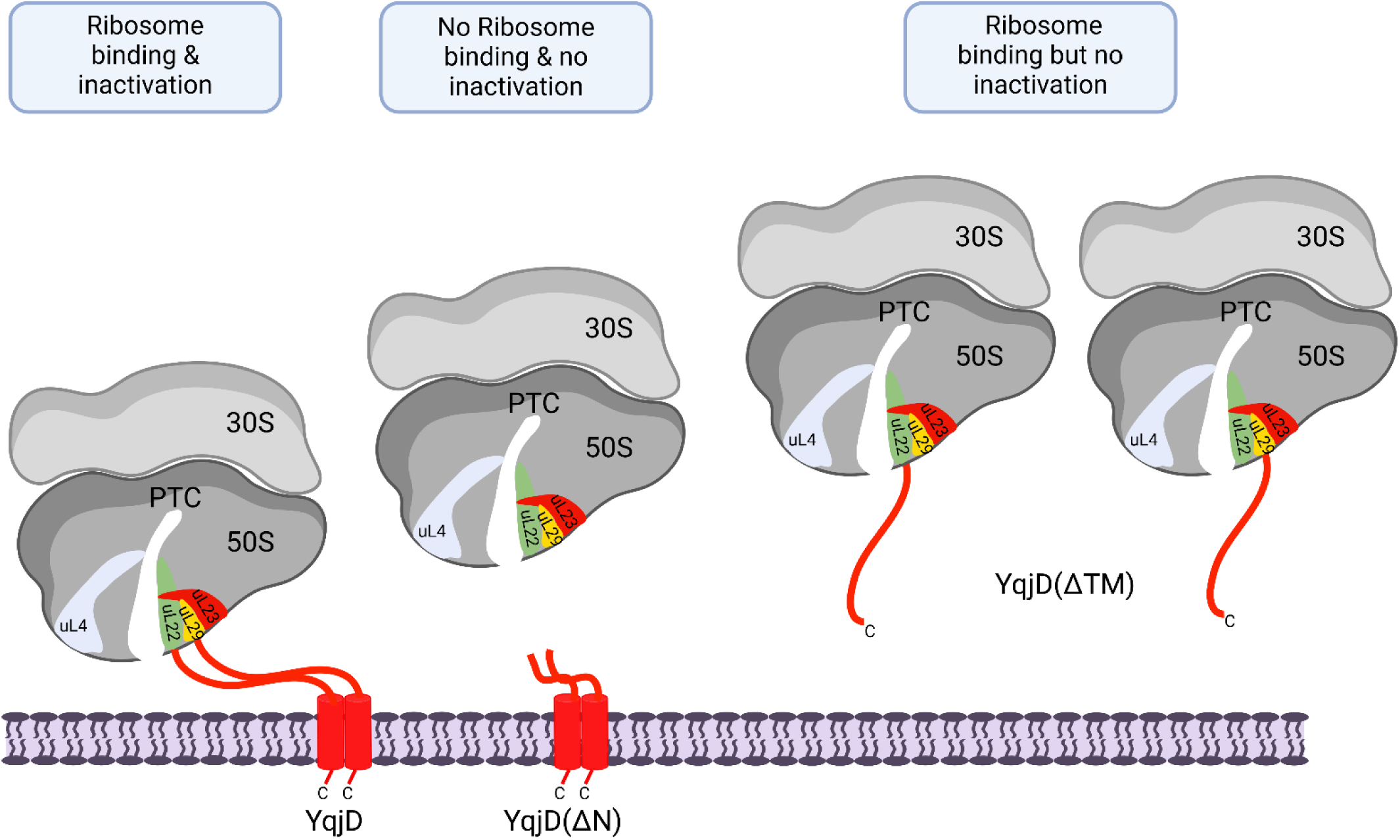
Hypothetical model for ribosome inactivation by YqjD. The C-tail anchored membrane protein YqjD shows a strong propensity for dimerization and interacts via its N-terminus with the ribosomal proteins uL22, uL23, and uL29. In addition, YqjD contacts uL24 (not shown). These proteins encircle the ribosomal peptide tunnel exit and form an important platform for ribosome-interacting proteins. As a consequence of YqjD binding to ribosomes, ribosomes are inactivated. N-terminally truncated YqjD variants fail to bind to ribosomes and are unable to prevent translation. Deletion of the C-terminal transmembrane domains prevents YqjD dimerization and ribosome inactivation, although this ΔTM-YqjD variant still shows ribosome binding, Thus, it is possible that for full ribosome inactivation, both N-termini of the YqjD dimer need to simultaneously bind to a single ribosome, but this needs to be further validated. Note, that the YqjD paralog, ElaB, can even protrude into the ribosomal tunnel.

C-tail anchored membrane proteins such as YqjD/ElaB/YgaM are generally rather rare in *E. coli* ^46^, and the benefit of having membrane-anchored hibernation factors in addition to several soluble hibernation factors is not entirely clear ^18,20,22^. YqjD and its paralogs are primarily located at the cell poles ^44,102^, where a large majority of ribosomes is also located ^103–105^. Thus, ribosome hibernation is promoted by co-localizing YqjD and a large majority of ribosomes at the cell poles during stationary phase. The determinants for the polar localization of YqjD still need to be further explored, but it has been suggested that specific interactions between YqjD’s TM and phosphatidic acid-rich membrane clusters are involved ^102^. This would then explain why YqjD/ElaB/YgaM require a TM. However, it is also possible that the oligomerization combined with ribosome binding, is sufficient to restrict YqjD’s diffusion in the membrane and to tether it to the cell poles. The polar localization also explains why moderate overexpression of *yqjD* is tolerated without drastic growth defects, because only those ribosomes located at the cell poles would be inhibited.

In conclusion, the membrane-localization of YqjD/ElaB/YgaM, their enrichment at the cell pole and their mechanism of ribosome inactivation by interacting with the ribosomal tunnel define them as a novel and widely distributed class of bacterial ribosome-hibernation factors.

## Acknowledgments

HGK, FD and PH gratefully acknowledge support from the Deutsche Forschungsgemeinschaft (DFG) (grants KO2184/8, KO2184/9 (SPP2002), and RTG 2202, Project-ID 278002225 to HGK, and SFB1381, Project-ID 403222702 to HGK, FD and PH). Work at the Molecular Foundry was supported by the Office of Science, Office of Basic Energy Sciences, of the U.S. Department of Energy under Contract No. DE-AC02-05CH11231. The work conducted at the U.S. Department of Energy Joint Genome Institute (https://ror.org/04xm1d337), a DOE Office of Science User Facility, is supported by the Office of Science of the U.S. Department of Energy operated under Contract No. DE-AC02-05CH11231.

## Author contributions

Conceptualization: HGK. Investigation: RN, JB, KS, EM, FD, CBH; Visualization: RN, JB, FD, CBH, HGK; Funding acquisition: HGK, CBH, FD, PH; Supervision: HGK; Writing: RN, JB, KS, EM, CBH, FD, PH, HGK. All authors have read and commented on the manuscript.

## STAR Methods

### RESOURCE AVAILABILITY

#### Lead contact

Further information and requests for resources and reagents should be directed to the lead contact, Hans-Georg Koch

##### Materials availability

All plasmids are available upon request, subject to a material transfer agreement (MTA), from Hans-Georg Koch

##### Data and Code availability

- All data reported in this paper will be shared by the lead contact upon request.
- The mass spectrometry proteomics data have been deposited to the ProteomeXchange Consortium via the PRIDE ^106^ partner repository with the dataset identifier PXD052307 and 10.6019/PXD05230. Any additional information required to reanalyze the data reported in this paper is available from the lead contact upon request.

##### Experimental model and subject details

All bacterial strains used in this study are derived from wild type *E. coli* K-12 strain 107-110. *E. coli* strains BW25113 or MC4100 served as wild-type strains and were routinely grown on LB medium 111 unless stated otherwise. The Δ*yqjD* (JW3069), Δ*elaB* (JW2261) and Δ*ygaM* (JW2647) strains were obtained from the Keio collection and were purchased via Horizon Discovery Ltd. (Cambridge, UK). Plasmids were propagated in *E. coli* DH5α 107. Proteins were expressed in *E. coli* strains BW25113, BL21, BL21(DE3) or C43(DE3) (Novagen/Merck, Darmstadt, Germany) for purification. HiFi DNA assembly Master Mix.

The sequence of all plasmids was confirmed by sequencing. All primers used in this study are listed in Supplementary Table S2.

## METHOD DETAILS

### Construction of YqjD, ElaB, and YgaM encoding plasmids

Plasmids encoding *yqjD, elaB* or *ygaM* with an N-terminal His-tag in a pBAD24 backbone (pBADHisA) were obtained from BioCat GmbH (Heidelberg, Germany). The YqjD variants were generated by using the Q5 site-directed mutagenesis kit (NEB, Frankfurt, Germany) and the following primer pairs. 1F/1R for the *ΔN(12)-*YqjD (deletion of the first 12 residues at the N-terminus region of YqjD), 2F/2R for the *ΔTM-*YqjD (deletion of the 18 residues (corresponding to the TM) at the C-terminus region of YqjD), 3F/3R for generation of YqjD(P_80_A) (the highly conserved Proline residue at position 80 in YqjD substituted to Alanine) and 4F/4R for generating YqjD(P_80_G) (the highly conserved Proline residue at position 80 in YqjD substituted to Glycine). The construction of the YqjD amber-stop codon variants for site-directed cross-linking used the primer pair 5F/5R for YqjD(L10pBpa) and 6F/6R for YqjD(I39pBpa). Additional YqjD truncations were generated and included the *ΔN(30)-*YqjD (deletion of the first 30-residues at the N-terminus region of YqjD) using primer pair 7F/7R, primers 8F/8R for *ΔN(45)-*YqjD (deletion of the first 45-residues at the N-terminus region of YqjD), 9F/9R for *ΔN(60)-*YqjD (deletion of the first 60-residues at the N-terminus region of YqjD). Deletion of the 6xHis and the Xpress-epitope tags attached to the N-terminus region of YqjD were deleted using primer pair 10F/10R. The nucleotide sequences of all primers is listed in Supp. Table S2. YqjD variants in which the nucleotide sequence encoding the YqjD-TM motif was substituted with the nucleotide sequence of the TM-domains of YfgM, FtsQ, or Fis1 were obtained from BioCat GmbH (Heidelberg, Germany). BioCat GmbH (Heidelberg, Germany) also synthesized the YqjD-TMD peptide.

### Viable cell staining and counting

The assay was performed using the QUANTOM Tx^TM^ Microbial Cell Counter and the QUANTOM^TM^ Viable Cell Staining Kit obtained from BioCat GmbH (Heidelberg, Germany) The kit stains live bacterial cells to be counted. The optical density of the cell-culture was determined using a spectrophotometer at each time point and an aliquot corresponding to approx. 1 x 10^8^ cells were collected. The cells were washed, the culture media was completely removed and the cells were resuspended in the QUANTOM^TM^ Viable Cell Dilution Buffer. 10.0 µL of the cell culture was taken into a clean 1.5mL Eppendorf-tube and 2.0µL of the QUANTOM^TM^ Viable Cell Staining Dye was added and mixed gently and carefully. The cells were then incubated at 37°C for 30 minutes in the dark. Thereafter 8.0µL QUANTOM^TM^ Cell Loading Buffer I was added and mixed gently without creating bubbles. 5.0µL of this mixture was loaded onto a QUANTOM^TM^ M50 Cell Counting Slide and centrifuged at 300 x rcf for 10 minutes in a QUANTOM^TM^ Centrifuge at room-temperature. The slide was then inserted into the QUANTOM Tx^TM^ cell counter and cells were counted with the light intensity level set to either 7 or 9. The obtained viable cell numbers were then blotted against time.

### Purification of YqjD, ElaB, YchF, SecA and SRP

The *E. coli* BW25113 strain expressing either ElaB, YqjD or its variants from the pBADHisA vector or *E. coli* BL21(DE3) expressing pET19b-SecA-His ^56^ or *E. coli ΔychF* (JW1194, Km^S^)^112^ expressing pBAD24-YchF-His were all grown on LB-medium supplemented with 100 µgmL^-1^ ampicillin to an optical density at 600 nm (OD_600_) of 0.5 and then induced with the corresponding inducers. For ElaB, YqjD and its mutants, as well as for YchF production, cells were induced with 0.02% arabinose for 1-2 hours, while for SecA production the cells were induced with 0.5 mM IPTG for 3-4 hours. Cell growth was stopped on ice for 10-15 minutes and cells were harvested at 7,460 x g for 15 minutes using a JLA-9.100 rotor (Beckman Coulter).

The cell pellet was resuspended in HKM buffer (25 mM HEPES, 200 mM KCl, 10 mM MgCl_2_ x 6 H_2_O and 10% glycerol, pH 7.5). Just before the cell lysis, 5 mM β-mercaptoethanol (β-ME), 0.5 mM PMSF (Carl Roth, Karlsruhe, Germany) and cOmplete^TM^ EDTA-free Protease Inhibitor Cocktail (1 tablet/ 50.0 mL cell culture, Sigma Aldrich, Germany) were added and the cell mixture homogenized using an IKA homogenizer (T 10 basic Ultra TURRAX^®^). Cells were then lysed by passing them 2-3 cycles through either Emulsiflex C3 (Avestin) or Maximator type HPL6 (Maximator GmbH) at 800-1000 bar (11,603 - 14,503 psi). The broken cells were cleared using a Sorvall RC6 (Thermo Scientific) at 30,000 x g for 30 minutes set at 4°C in an SS34 rotor and the cell debris discarded. The S30 (cleared cell lysate or supernatant) was further centrifuged at 183,700 x g for 2½ hours in a Ti50.2 rotor using Sorvall WX-90 Ultra Series (Thermo Scientific) set at 4°C. For membrane proteins, the supernatant (S150) was discarded, and the crude membrane pellet was homogenized using a Dounce homogenizer in solubilization buffer (HKM buffer plus 1.0% n-dodecyl-β-D-maltoside (DDM, Carl Roth), 5 mM β-ME, 0.5 mM PMSF and cOmplete^TM^ EDTA-free Protease Inhibitor Cocktail). Solubilization was performed for 1 h at 4 °C. The solubilized membranes were centrifuged using a Sorvall RC6 centrifuge (Thermo Scientific) at 30,000 x g for 15 minutes at 4°C in an SS34 rotor. The solubilized materials were added directly to the equilibrated TALON^®^ resin (TaKaRa; Clonetech) and incubated for 2.0 hours on a rotating shaker. For cytosolic proteins, the S150 (supernatant after the ultracentrifugation at 183,700 x g for 2½ hours in a Ti50.2 rotor at 4°C) was directly added to the equilibrated TALON^®^ resin and incubated for 2.0 hours on a rotating shaker. The resin was washed 3-times, for 15 minutes each with washing buffer (HKM buffer with 0.03% DDM and 5 mM imidazole pH 8.0). The washed TALON^®^ material was centrifuged using an Eppendorf 5804R centrifuge at 2,937 x g for 5 min at 4°C. The TALON^®^ resin were then transferred into 15 mL polypropylene columns and proteins were eluted with elution buffer (HKM buffer with 0.03% DDM) containing initially 20 mM imidazole pH 8.0 and then 200 mM imidazole. The eluted proteins were buffer-exchanged using PD-10 desalting columns Disposable (Sigma Aldrich; Merck) against storage buffer (50 mM HEPES, 50 mM potassium acetate, 10 mM magnesium-acetate and 1.0 mM DTT pH 7.5). The buffer was supplemented with 0.03% DDM if membrane proteins were to be stored. The concentrations were determined by either BCA assays or A_280_, while the purity was established on an SDS-PAGE. The proteins were aliquoted, frozen in liquid nitrogen and kept at −80 °C for further usage.

For the purification of SRP, its protein component Ffh was expressed from pTrc99a-His-Ffh in *E. coli* BL21 ^113^. Cells were induced at an OD_600_ of 0.5 with 1 mM IPTG for 3 h, harvested, washed and lysed with the Emulsiflex Homogenizer (Avestin Europe, Mannheim, Germany). The lysate was cleared at 30,000 x g for 20 min at 4 °C in an SS-34 rotor and loaded on buffer-equilibrated (25 mM HEPES-KOH, 1 M NH-acetate, 10 mM Mg-acetate, 1 mM β-mercaptoethanol, 15% glycerol, 5 mM imidazole, pH 7.6) Talon beads for 1 h. After several washing steps with the equilibration buffer, Ffh was eluted with the same buffer containing 200 mM imidazole, re-buffered into HT buffer (50 mM HEPES/KOH pH 7.5, 100 mM potassium acetate pH 7.5, 10 mM magnesium acetate pH 7.5, 1 mM DTT, 50% glycerol) and stored at - 20 °C. A reconstitution of Ffh with 4.5S RNA is usually not required, as Ffh has sufficient 4.5S RNA bound ^113,114^.

### Ribosome and membrane purification

High-salt washed wild-type 70S ribosomes were purified from strains MC4100 or BW25113. Ribosomes bearing pBpa at uL23 were purified from MC4100*ΔrplW::kan* containing pCDF-L23(E52pBpa) or L23(G71pBpa) ^66^ and pSup-BpaRS-6TRN ^63^ for pBpa incorporation. The cells were propagated in S150-medium consisting of 1% (g/w) yeast extract, 1% (g/w) tryptone-peptone, 41 mM KH_2_PO_4_, 166 mM K_2_HPO_4_ and 1% (g/w) glucose. Except for wt ribosomes, the medium was supplemented with 50 μg/mL streptomycin (Sigma Aldrich) and 0.5 mM IPTG. For pBpa incorporation, 35 μg/mL chloramphenicol (Sigma Aldrich) and 0.5 mM pBpa (Bachem, Bubendorf, Switzerland) were added to the medium. When the cell density reached OD_600_ of 1.6-1.8, the growth was stopped on ice for 10-15 min, and the cells were harvested, washed and homogenized with the Emulsiflex C3. The lysate was cleared at 30,000 x g for 30 min and the crude ribosomes were collected at 184,000 x g in a Ti50.2 rotor for 2.5 h. Ribosomes were dissolved in high-salt buffer (50 mM Triethanolamine acetate; 1M potassium acetate; 15 mM magnesium acetate; 1 mM DTT; pH 7.5) and purified through a 1.44 M sucrose cushion at 344,000 x g for 1 h. 70S ribosomes were isolated via centrifugation through a 0.29-1.17 M sucrose gradient at 29,000 rpm for 17 h in a TH-641 swing-out rotor (Thermo Fisher Scientific). The ribosomal fractions were withdrawn from the gradient, concentrated at 344,000 x g for 1h in a TLA120.2 rotor and resuspended in CTF-buffer at pH 7.5 with 1 mM DTT. The isolation of ribosomes from stationary phase *E. coli* cells followed the same protocol, but *E. coli* cells were grown for 24 h up to an optical density of about 6.0.

For membrane isolation, cells were grown to approx. OD_600_= 1.5-1.8 on LB medium, harvested and resuspended in INV buffer (50 mM triethanolamine acetate, pH 7.5, 250 mM sucrose, 1 mM EDTA, 1 mM DTT) supplemented with 0.5 mM PMSF and cOmplete^TM^ EDTA-free Protease Inhibitor Cocktail. Next, the samples were lysed as described above and the cell debris was removed by centrifugation at 30,000 x g for 30 minutes in an SS34 rotor. The supernatant (S30) was further centrifuged at 184,000 x g for 2,5 hours at 4 °C in a Ti50.2 rotor and the pellet containing the crude bacterial membranes was dissolved in INV buffer, loaded onto a 10-30% sucrose gradient and the inner membrane fraction (inverted inner membrane vesicles, INV) and the outer membrane fraction were separated as described ^115^.

### Ribosome binding assays

For ribosome-YqjD binding assays, 10 nM of the purified wild-type YqjD was incubated with varying concentrations of sucrose-gradient purified *E. coli* ribosomes in 50 mM HEPES, 50 mM potassium acetate, 10 mM magnesium-acetate, pH 7.5, 1.0 mM DTT, 0.03% DDM and 1.0 mM spermidine. Ribosomes were isolated as described above from cells grown to either exponential (2.5 h) or stationary phase (24 h) and prepared in high salt buffer. After mixing the components, they were incubated at 30 °C for 30 minutes. The reaction mix was overlaid onto a 30% sucrose cushion (in the above buffer) in TLA 120.2 rotor tubes and centrifuged at 344,000 x g for 2.0 hours. The supernatant was carefully collected into a separate tube, while the pellet was also resuspended and collected in a second tube. The proteins were precipitated by adding 10% TCA, denatured and separated by SDS-PAGE, followed by western blotting.

### *In vitro* protein synthesis

For *in vitro* protein synthesis, a purified transcription/translation system composed of cytosolic translation factors (CTF) and high salt washed ribosomes ^55^ was used. The ^35^S-Methionine/ ^35^S-Cysteine labeling mix was obtained from Hartmann Analytics (Braunschweig, Germany). After 30 min at 37 °C, the *in vitro* reaction was directly precipitated with 10% trichloroacetic acid (TCA). Next, the samples were denatured at 56 °C for 10 minutes in 25 μl of TCA loading dye (prepared by mixing one part of Solution III (1M dithiothreitol) with 4 parts of Solution II (8.3% SDS (w/v), 0.083 M Tris-Base, 30% glycerol and 0.03% Bromophenol blue) and 5 parts of Solution I (0.2 M Tris, 0.02 M EDTA pH 8)) and analyzed on SDS-PAGE and by phosphor imaging.

### Immune detection and antibodies

For immune detection after SDS-PAGE, samples were electro blotted onto Nitrocellulose 0.45 µm membranes (GE Healthcare) with a current of 750 mA for 2.5 hours in a tank buffer system (transfer-buffer: 20 mM Tris, 150 mM Glycine, 20% Ethanol (v/v), 0.02% SDS (w/v)). Membranes were blocked with 5% milk powder in T-TBS buffer for at least 1 h. Polyclonal antibodies against YidC and Ffh were raised in rabbits against the complete and SDS-denatured protein ^55,116^. Monoclonal antibodies against the His6-tag were purchased from Thermo Scientific and from Roche. Antibodies against the Xpress epitope tag were purchased from Invitrogen Life technologies. Peroxidase-coupled goat anti-rabbit and goat-anti mouse antibodies from SeraCare were purchased via medac GmbH (Wedel, Germany) and were used as secondary antibodies with ECL (GE Healthcare) ^55,117^. Antibodies against *E. coli* ribosomal proteins were raised in sheep and were a gift from Richard Brimacombe (Max-Planck-Institut für Molekulare Genetik, Berlin).

Peptide antibodies against YqjD (MSKEHTTEHLRAEL), ElaB (VLRSSGDP-ADQKYV) and YgaM (GSDAKGEAEAARSK) were raised in rabbits by GeneScript (Leiden, Netherlands).

### *In vivo* and *in vitro* cross-linking

p-benzoyl-l-phenylalanine (pBpa) for cross-linking was obtained from Bachem (Bubendorf, Switzerland). For site-directed *in vivo* cross-linking C43(DE3) cells containing the amber-stop codon variants of YqjD on a plasmid and pEVol were cultured overnight in LB medium at 37 °C. 10 ml of the overnight culture were used for inoculation of 1000 ml LB medium supplemented with 1 ml pBpa (final concentration 0.5 mM, dissolved in 1 M NaOH), 100 μg/μl of ampicillin and 25 μg/μl of chloramphenicol. The cultures were further incubated at 37 °C until they reached the early exponential growth phase (OD_600_ = 0.5-0.8) and induced with 0.02% arabinose. After induction, the cultures were grown for 1-2 hours at 37 °C, cooled down on ice for 10-15 minutes and harvested by centrifugation at 7,460 x g in a JLA 9.1000 rotor for 15 minutes. The cell pellets were resuspended in PBS buffer (137 mM NaCl, 2.7 mM KCl, 10 mM Na_2_HPO_4_ and 1.8 mM KH_2_PO_4_), harvested again as above, resuspended in 10 ml PBS buffer, and divided into two multi-well plates. One plate was exposed to UV light (365nm) on ice for 30 minutes (UV chamber: BLX-365, from Vilber Lourmat) while the other plate was kept in the dark. After UV irradiation, the cell suspension was transferred to SS34 tubes and cells were collected by centrifugation at 11,950 x g for 15 minutes. Each cell pellet was resuspended in 10 ml of lysis buffer (25 mM HEPES, 200 mM KCl and 10 mM MgCl_2_ x 6H_2_O) and 10% glycerol, pH 7.5), including protease inhibitors (0.5 mM PMSF and cOmplete^TM^ EDTA-free Protease Inhibitor Cocktail) and YqjD was purified as described above.

For *in vivo* crosslinking with the uL23-pBpa variants, the strain MC4100*ΔrplW::kan* containing pCDF-L23(E52pBpa) or L23(G71pBpa) ^57,66^ and pSup-BpaRS-6TRN ^63^ was used and the LB medium additionally supplemented with 50 μg/mL streptomycin and 0.5 mM IPTG. Cells were either grown to OD_600_ = 1 (exponential phase) or OD_600_ = 4.5 (stationary phase), cooled, harvested, UV exposed and then lysed in CTF buffer (50 mM triethanolamine acetate pH 7.5, 50 mM potassium acetate pH 7.5, 5 mM magnesium acetate pH 7.5, 1 mM DTT) as described above. Subsequently, bacterial membranes were prepared, and crude ribosomes purified as before. 500 µg of total protein for each sample was TCA-precipitated, separated by SDS-PAGE and analyzed by Western blot.

For *in vitro* crosslinking, purified *E. coli* 70S ribosomes (500 nM) were combined with purified SRP (500 nM) or purified YqjD (5 µM), mixed 20 min at 30 °C in CTF buffer in a total volume of 50 µl and crosslinked by UV exposure using a Biolink 365 nM-crosslinking chamber (Vilber-Lourmat) for 30 min on ice. Samples were then TCA-precipitated, separated on PAGE, and analyzed by western blot.

### Bioinformatic analyses

DUF883-containing proteins were identified by searching the UniProt database ^118^ for the InterPro domain IPR010279 ^119^. The sequence similarity network was built using the EFI-EST webtool ^120^ using IPR010279 with an alignment score of 20. Nodes were collapsed at 95% identity. The network was visualized with Cytoscape v3.10.1 using the Prefuse Force Directed OpenCL Layout. Nodes were colored based on the phylum of the representative sequence. Phylogenetic trees were built using MAFFT for multiple sequence alignments on the CIPRES Science Gateway ^23,121^ with default parameters, and either the IQ-Tree web server ^122,123^ with LG+F+G4 (the best-fit model according to the Bayesian information criterion, BIC) ^124^ and ultrafast bootstrap (1000 replicates) ^125^ for the tree in Suppl. Figure S3 (and extracted subtrees in Fig. 2B), or with FastTreeMP on XSEDE ^126^ for the tree in Suppl. Fig. 2A. The resulting trees were visualized and annotated in iTol ^127^. Sequences for the tree in Suppl. Fig. S3A were selected based on the clusters identified in the SSN, such that each numbered cluster contained at least two representatives, and for any given organism, all paralogs were included. In this way, some clusters are represented by more than 2 sequences. For the tree in Suppl. Fig. S2A, YqjD, YgaM and ElaB were used to search UniRef90 clusters in the UniProt database with blastp; the top 250 UniRef90 clusters were collected and duplicates removed. The representative sequences for the resulting UniRef90 clusters were used to build the tree.

TM regions were predicted using DeepTMHMM ^128^. AlphaFold2-predicted structures were downloaded from the Alpha Fold Protein Structure Database ^129^. ChimeraX ^130^ was used to visualize structural models and perform conservation analysis using AL2CO ^131^.

### Mass spectrometry

For identification of UV-crosslinked proteins, gel lanes from SDS-PAGE of UV-illuminated or control samples analysis were cut in 26 slices which were individually subjected to in-gel protein digestion using trypsin as previously described ^11^. Peptide mixtures were desalted using self-packed C18 Stop And Go Extraction tips^132^ using two layers of 1.0 × 1.0 mm C18 Empore Disks (3M, St. Paul, USA) and directly analysed by liquid chromatography-tandem mass spectrometry (LC-MS/MS). For chromatographic separation an Ultimate 3000 RSLCnano system was coupled online to an Orbitrap Elite mass spectrometer (Thermo Fisher Scientific, Bremen, Germany). Peptides were washed and preconcentrated on nanoEase™ M/Z Symmetry C18 pre-columns (20 mm x 180 μm inner diameter; Waters) at a flow rate of 10 µl/min and separated using nanoEase™ M/Z HSS C18 T3 columns (25 cm x 75 μm inner diameter; pore size, 100 Å; particle size, 1.8 μm; Waters) at a flow rate of 0.3 µl/min and 40°C. Peptide elution was controlled with a binary solvent system consisting of 0.1% (v/v) FA (solvent A) and 0.1% (v/v) FA/50% (v/v) MeOH/30% (v/v) ACN (solvent B) using a gradient of 7 - 65% solvent B in 30 min, 65 - 80% in 5 min, and 3 min at 80%, interfaced online with the Nanospray Flex ion source with PST-HV-NFU liquid junction (MS Wil, The Netherlands) and fused silica emitters (EM-20-360; MicrOmics Technologies LLC). MS instruments were operated in data-dependent mode with parameters as follows: mass range of *m/z* 370 - 1,700 for MS1 scans with a resolution of 120,000 (at *m/z* 400), a target value (AGC) of 1 x 10^6^ ions, and a maximum injection time (IT) of 200 ms. For MS2 scans, up to 15 most intense precursor peptides with a charge ≥ 2 were selected for collision induced dissociation (CID) in the linear trap with a normalized collision energy of 35%, activation time 10 sec, q-value=0.25, a resolution of 35,000, AGC of 5 x 10^4^ ions, a max. IT of 150 ms, and dynamic exclusion for 45s. Mass spectrometric raw data were processed using MaxQuant (version 2.0.2.0; ^133^) searching against the *E.coli*-specific database from UniProt (release 2022_01). Database searches were performed with tryptic specificity and a maximum number of two missed cleavages. Mass tolerances were set to 4.5 ppm for precursor ions and 0.5 Da for fragment ion matchings. Carbamidomethylation of cysteine residues was considered as fixed modification, oxidation of methionine and N-terminal protein acetylation were set as variable modifications. The options ‘match between runs’ and ‘iBAQ’ were enabled. Proteins were identified with at least one unique peptide and a false discovery rate of 0.01 on both peptide and protein level.

### Data quantification and statistical analyses

Western blot and autoradiography samples were analyzed by using the *ImageQuant* (GE Healthcare) or *ImageJ/ Fiji* plug-in software (NIH, Bethesda, USA). All experiments were performed at least three-times as independent biological replicates and representative gels/blots/images are shown. When data were quantified, at least three independent biological replicates with several technical replicates were performed. Mean values and SEM values were determined by using either Excel (Microsoft Corp.) or GraphPad Prism (GraphPad Prism Corp. San Diego).

## Supplemental information

Table S1: Excel file containing additional data too large to fit in a PDF, related to the YqjD sequence similarity network nodes in Figs 1, 2, S1, S2.

Document S1: Table S2 and Figures S1-S7, related to Figs. 1-8.

