## Supplemental information for "Ribosome-inactivation by a class of widely distributed C-tail anchored membrane proteins"

### Supplemental Table 2: Oligonucleotide sequences used for PCR reactions

| Primer Name | | | Primer sequence | Modification |
|---|---|---|---|---|
| 1 | Fwd Primer | 1F | 5′-gagttgaaatccctttccgatacg-3′ | *Δ12AA-N-term*YqjD<br>Deletion of the first 12 residues at the N-terminus region of yqjD |
| | Rev Primer | 1R | 5′-ctcgagctcggatcccca-3′ | |
| 2 | Fwd Primer | 2F | 5′-tcgcgtcgttaaaagcttg-3′ | *Δ18AA-C-term*YqjD<br>Deletion of 18 residues at the C-terminus region of YqjD |
| | Rev Primer | 2R | 5′-cggattttcacgcacatac-3′ | |
| 3 | Fwd Primer | 3F | 5′-gcgtgaaaatgcgtggacgggcg-3′ | YqjD$_{P80A}$<br>Substitution of Proline residue with Alanine |
| | Rev Primer | 3R | 5′-acatactcatcggcacgcgc-3′ | |
| 4 | Fwd Primer | 4F | 5′-gcgtgaaaatggctggacgggcgtgg-3′ | YqjD$_{P80G}$<br>Substitution of Proline residue with Glycine |
| | Rev Primer | 4R | 5′-acatactcatcggcacgc-3′ | |
| 5 | Fwd Primer | 5F | 5′-tacggaacattagcgtgctgagt-3′ | YqjD-L10pBpa<br>Substitution of leucine (ctg) at position 10 with amber stop codon (tag) |
| | Rev Primer | 5R | 5′-gtgtgttctttcgacatctc-3′ | |
| 6 | Fwd Primer | 6F | 5′-gttgagtaagtagcgtagcaaagc-3′ | YqjD-I39pBpa<br>Substitution of isoleucine (ttg) at position 39 with amber stop codon (tag) |
| | Rev Primer | 6R | 5′-tcttcttttgacttctcg-3′ | |
| 7 | Fwd Primer | 7F | 5′-aagtcaaaagaagagttgagtaagattcg-3′ | *Δ30AA-N-term*YqjD<br>Deletion of the first 30 residues at the N-terminus region of yqjD |
| | Rev Primer | 7R | 5′-ctcgagctcggatcccca-3′ | |
| 8 | Fwd Primer | 8F | 5′-gcactgaaacagagccgttatc-3′ | *Δ45AA-N-term*YqjD<br>Deletion of the first 45 residues at the N-terminus region of yqjD |
| | Rev Primer | 8R | 5′-ctcgagctcggatcccca-3′ | |
| 9 | Fwd Primer | 9F | 5′-attgccaaacaaacccgtgtc-3′ | *Δ60AA-N-term*YqjD<br>Deletion of the first 60 residues at the N-terminus region of yqjD |
| | Rev Primer | 9R | 5′-ctcgagctcggatcccca-3′ | |
| 10 | Fwd Primer | 10F | 5′-atgtcgaaagaacacactac-3′ | Deletion of the His-tag and the Xpress-epitope tag added/ linked to the N-terminus region of yqjD |
| | Rev Primer | 10R | 5′-agaaccccccatggttaa-3′ | |

**Legends to Supplementary Figures.**

**Figure S1: Domain structure of YqjD, ElaB and YqaM.** Related to Figure 1. Screenshots of YqjD, ElaB, and YgaM domain architectures in the InterPro database. The primary amino acid sequence and corresponding coordinates are shown at the top of each schematic followed by a cartoon of the confidence values associated with the precomputed AlphaFold structures (shown in Fig. 1B) and matching models for domains, families and other features. The interactive schematics can be viewed at the InterPro website:

YqjD: https://www.ebi.ac.uk/interpro/protein/UniProt/P64581/ ,

ElaB: https://www.ebi.ac.uk/interpro/protein/UniProt/P0AEH5/ ,

YgaM: https://www.ebi.ac.uk/interpro/protein/UniProt/P0ADQ7/ .

**Figure S2: Evolution of the YqjD, ElaB and YgaM subfamilies.** Related to Figures 1 and 2. (A) Phylogenetic tree of YqjD, ElaB and YgaM and closely related homologs. The tree was rooted at the clade containing proteins from SSN clusters 1 and 4 (Fig. 1C). Clades containing the YqjD, YgaM, and ElaB subfamilies are shaded and labeled. The proteins within each colored clade were used for the subfamily conservation analyses. (B) Sequence logos representing the multiple sequence alignments containing proteins from each subfamily. The TM regions predicted with DeepTMHMM are shown with the green box.

**Figure S3: YqjD, ElaB and YgaM represent three distinct subfamilies conserved in Enterobacterales.** Related to Figures 1 and 2. (A) Phylogenetic reconstruction of representative DUF883-containing proteins under maximum likelihood. The "SSN cluster" column corresponds to clusters 1-10 in Fig. 1C. Taxonomic classification for each node is given according to the color key. The protein labels for each leaf list the organism, followed by the cluster number corresponding to Fig. 1C, followed by the paralog number. If the paralog number is "0', then that organism has a single DUF883-containing protein, where as "1" indicates 1 of N paralogs. Bootstrap values greater than 0.5 are represented with a normalized purple circle according to the key. The YqjD, ElaB and YgaM clades are shaded with a light blue. (B) For each protein in panel A, the gene neighborhood is given. Genes are colored according to shared domains; light pink genes encode proteins with no identified domain. All genes are shown as transparent, except for those genes and homologs listed in the key. (C) Protein abundance of YqjD, ElaB and YgaM in *E. coli* BW25113 grown on M9 medium + 5g/l glucose to either exponential phase or to stationary phase. Data were extracted from (Schmidt *et al.*, 2016). IBAQ; Intensity Based Absolute Quantification.

**Figures S4: Ribosome binding of YqjD.** Related to Figure 4. (A) Purified YqjD (10 nM) was incubated at 30 °C with increasing concentrations of sucrose gradient-purified ribosomes, which were isolated from *E. coli* cells grown to either exponential (2.5 h) or stationary phase (24 h). After incubation for 30 min, the samples were centrifuged though a sucrose cushion and the amount of the YqjD in the pellet and in the supernatant were analyzed by immune detection. For quantification, the signal intensities of both supernatant and pellet were set to 100% and the percentage of ribosome-bound YqjD in the pellet fraction was quantified using *ImageJ*. (B) Wt, wt cells containing pBAD-YqjD and Δ*yqjD* cells were grown overnight on LB medium, sequentially diluted in PBS and 20 µl of each dilution was spotted on LB-plates and LB-plates containing low concentrations of chloramphenicol. Note that the panels from wt, wt + pBAD-YqjD and Δ*yqjD* cells were derived from the same plate, but unrelated areas of the plate were removed.

**Figures S5: ElaB inhibits *in vitro* protein synthesis.** Related to Figure 4. (A) ElaB was purified via affinity chromatography and added to a cell-free *E. coli in vitro* transcription/translation system (Koch *et al.*, 1999), synthesizing the $^{35}$S-labelled inner membrane protein mannitol permease (MtlA) as described in the legend to Fig. 4. Samples were separated by SDS-PAGE and analyzed by phosphorimaging. (B) Quantification of *in vitro* MtlA synthesis in the presence of purified ElaB. Shown are the mean values and the error bars represent the SD. The YqjD data shown in Figure 4D are displayed again as a reference. (C) *E. coli* cells expressing YqjD, ElaB or YgaM were TCA-precipitated, denatured at 56 °C and analyzed with α-Xpress antibodies after separation by SDS-PAGE and western blotting. Indicated are monomeric YqjD, ElaB and YgaM, and dimeric YqjD. The band labeled with * likely represents a degradation product of the YqjD dimer.

**Figure S6: The role of YqjD's transmembrane domain in ribosome inactivation.** Related to Figure 5. (A) Purified YqjD, YqjD(ΔN12) or purified YqjD(ΔTM) (10 nM each) were incubated at 30 °C with increasing concentrations of sucrose gradient-purified ribosomes, which were isolated from *E. coli* cells grown to exponential  phase (2.5 h). After incubation for 30 min, the samples were centrifuged though a sucrose cushion and the amount of the YqjD variants in the pellet and in the supernatant were analyzed by immune detection. For quantification, the signal intensities of both supernatant and pellet were set to 100% and the percentage of ribosome-bound YqjD in the pellet fraction was quantified using *ImageJ*. The values represent the means of the three independent experiments and the error bar reflects the SD. (B) *E. coli* wild type cells (BW25113) expressing YqjD with (pBad24-YqjD) or without His-tag (pBad24-YqjD$_{\Delta\text{His}}$) were separated on SDS-PAGE and after western blotting analyzed by α-YqjD peptide antibodies. Cells without plasmid and antibodies against YidC were used as a control. (C) *E coli* cells expressing YqjD variants that contained the native transmembrane domain (wt TM) or the transmembrane domain of YfgM, Fis1 or FtsQ were purified and tested for the ability to reduce in vitro protein synthesis of MtlA *in vitro*. MtlA synthesis was analyzed by phosphorimaging and MtlA synthesis in the absence of YqjD/YqjD variants was set as 100%. Shown are the mean values of three independent experiments and the error bar represents the SD. (D) Purified YqjD or its variants with different TMs were TCA precipitated, denatured at 56 °C and analyzed by immune detection with α-Xpress antibodies.

**Figure S7: YqjD interacts with proteins surrounding the ribosomal tunnel exit.** Related to Figures 6 and 7. (A) Samples were processed as in Fig. 7 and affinity purified YqjD and its cross-linked partner proteins were separated on SDS-PAGE. The gel lanes were subsequently sliced into multiple slices, which were separately processed and analyzed by mass spectrometry. Shown are the normalized intensities of uL24 and uL23 peptides in the –UV and +UV treated samples of wild-type YqjD, YqjD(I39) and YqjD(L10). (B) The *E. coli* Δ*rplW* (uL23) deletion strain expressing uL23(E52pBpa) or uL23(E71pBpa) was grown to exponential or stationary phase and UV-exposed. Subsequently, cells were fractionated into the soluble ribosome fraction and the crude membrane fraction. These fractions were then separated by SDS-PAGE and probed with antibodies against YqjD.

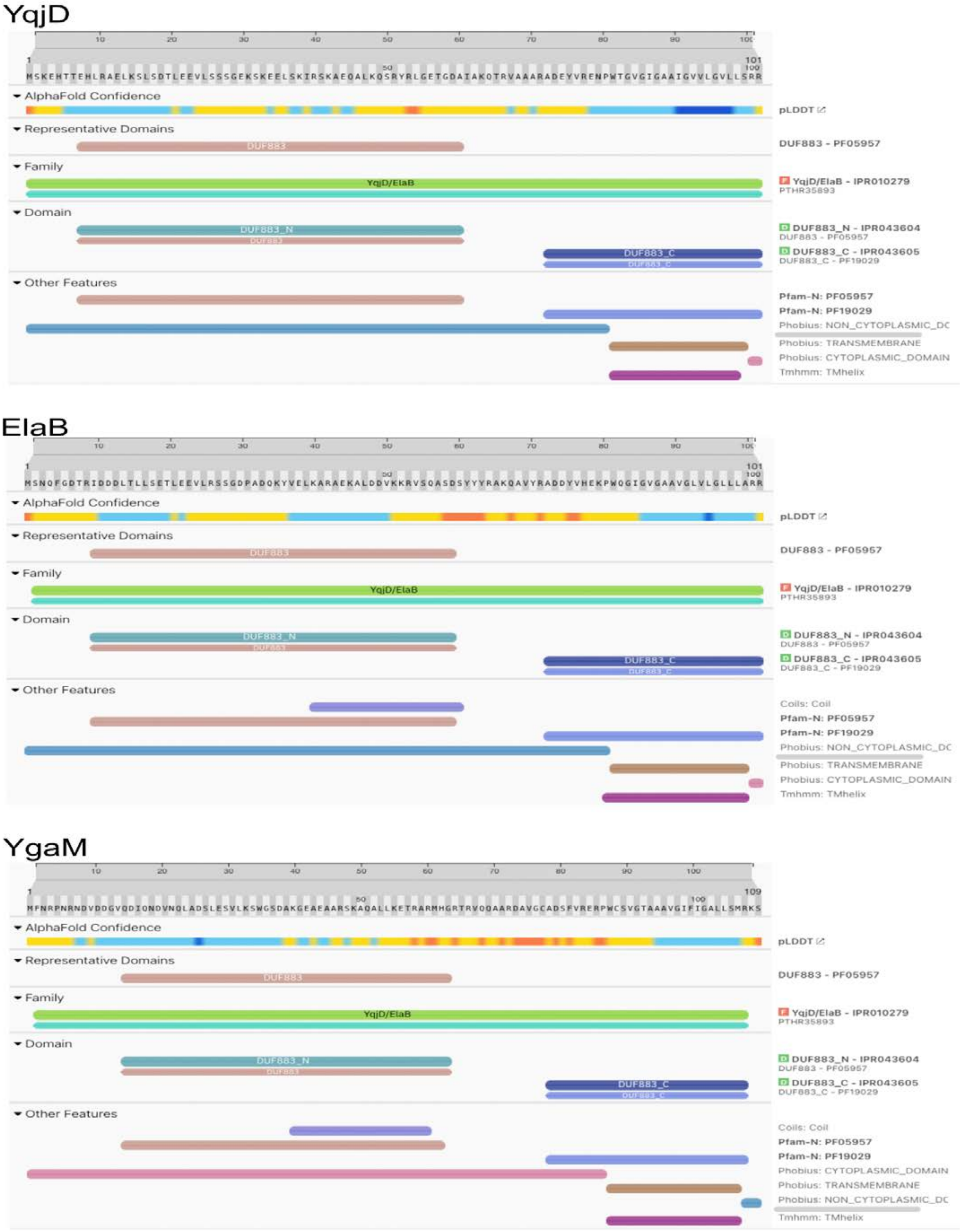


(Njenga et al., Supp. Fig. 1)

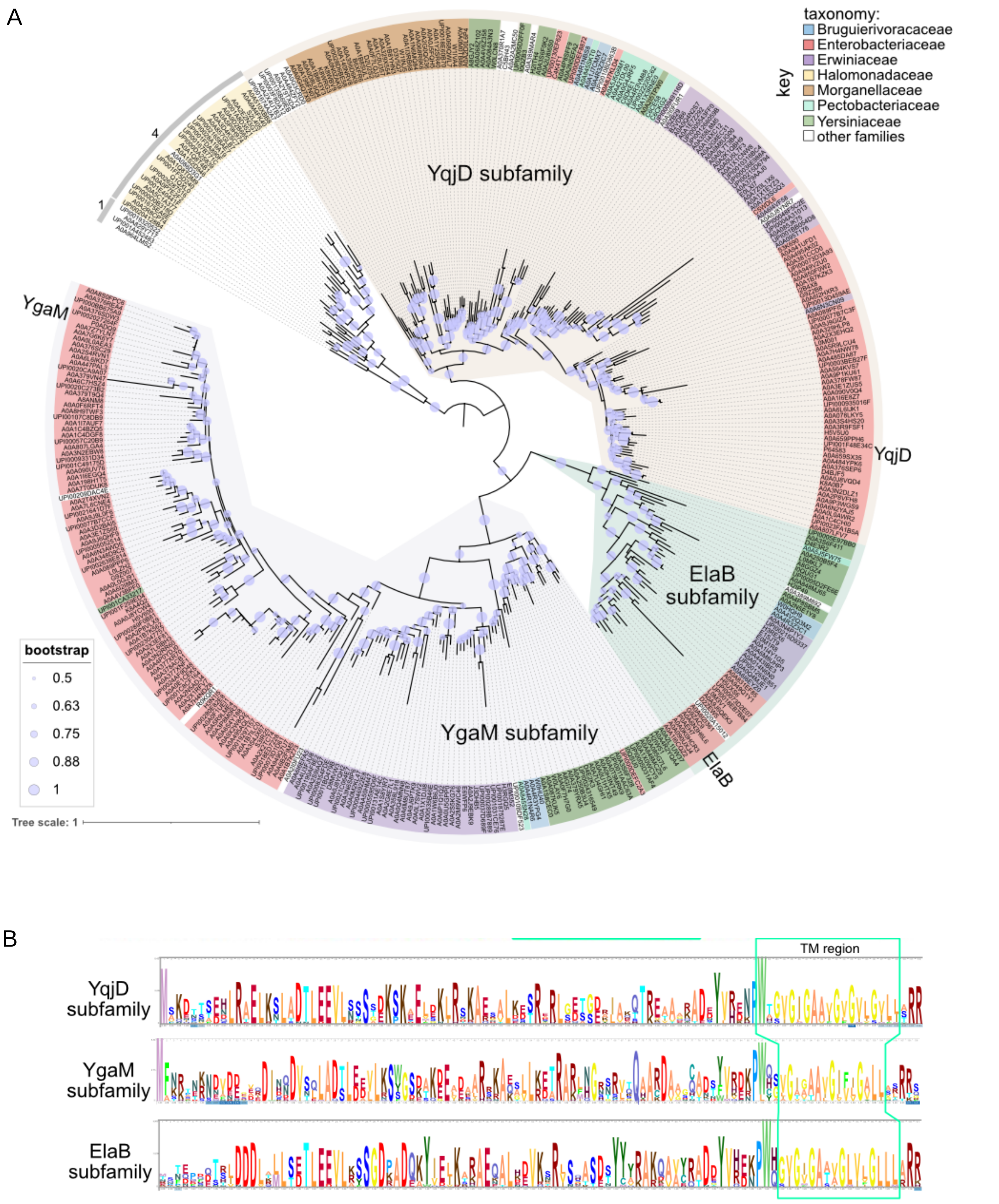


(Njenga et al., Supp. Fig. 2)

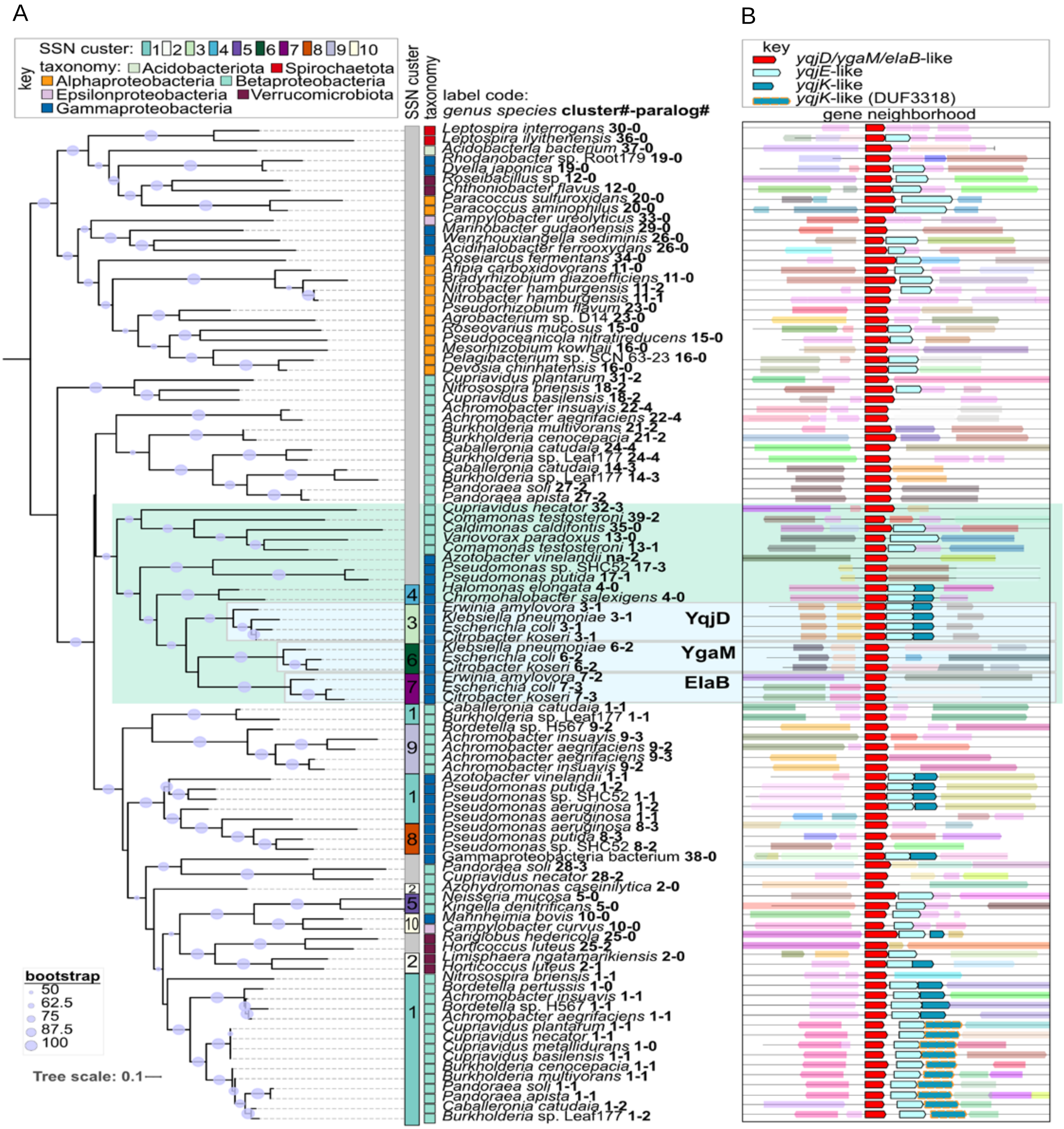


C

| Protein abundance (proteins/cell) | | |
|---|---|---|
| | IBAQ exponential phase | IBAQ stationary phase |
| YqjD | 2146 | 6209 |
| ElaB | 2398 | 4052 |
| YgaM | 762 | 793 |

(Njenga et al., Supp. Fig. 3)

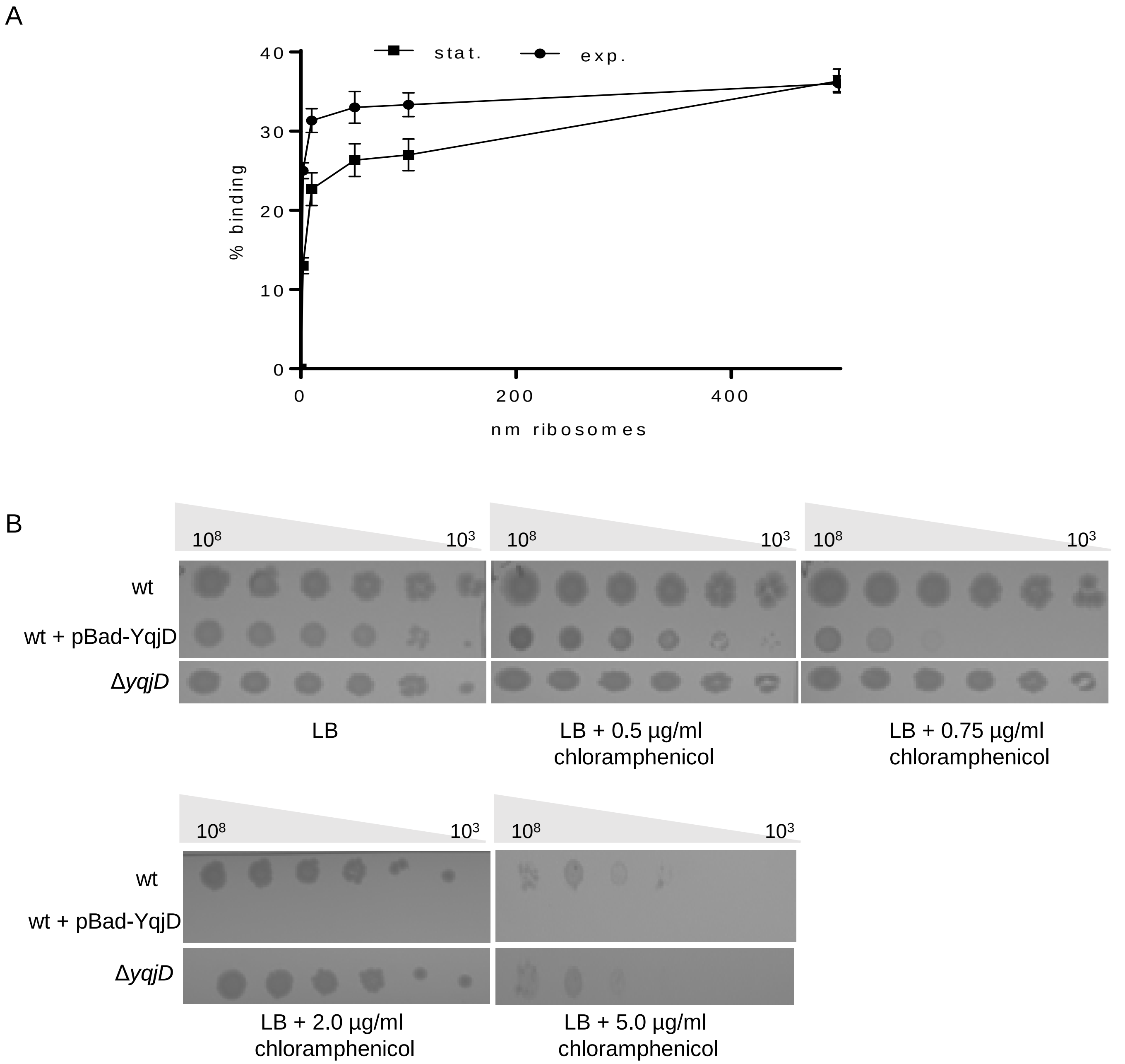


(Njenga et al., Supp. Fig. 4)

A

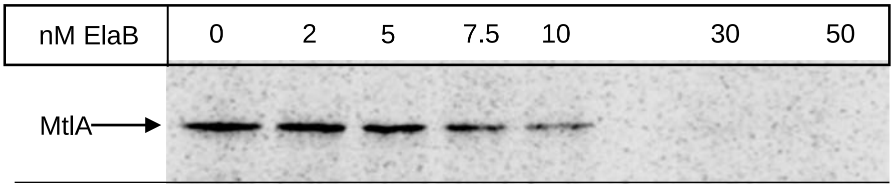


B

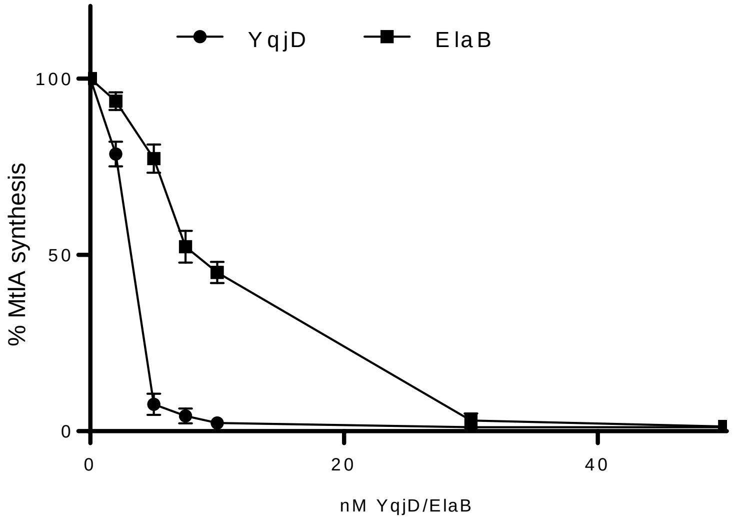


C

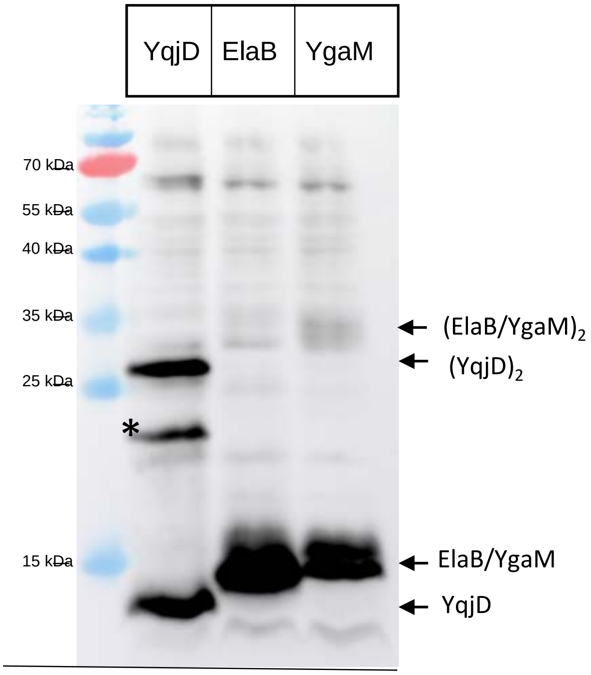


(Njenga et al., Supp. Fig. 5)

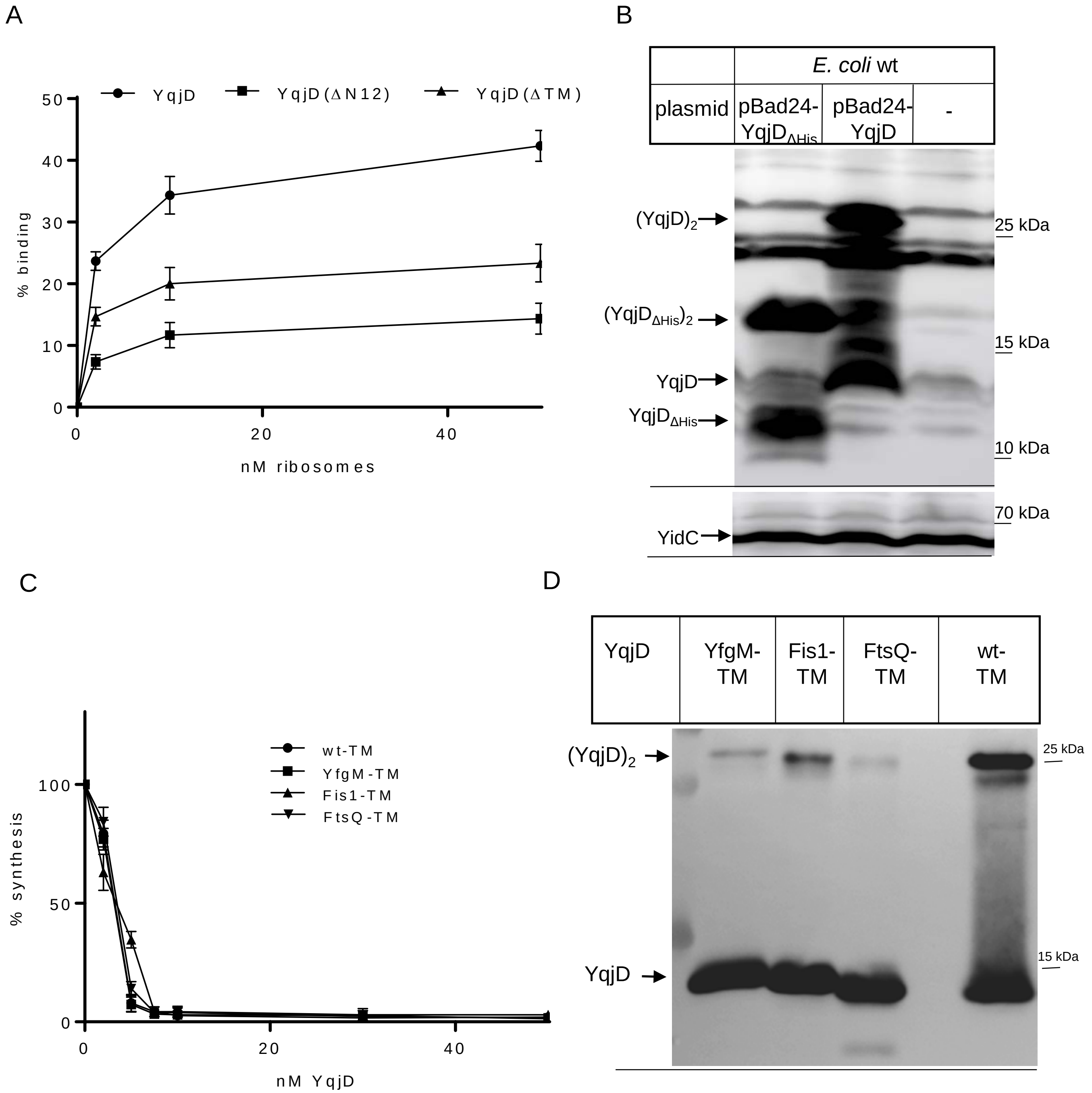


(Njenga et al., Supp. Fig. 6)

A

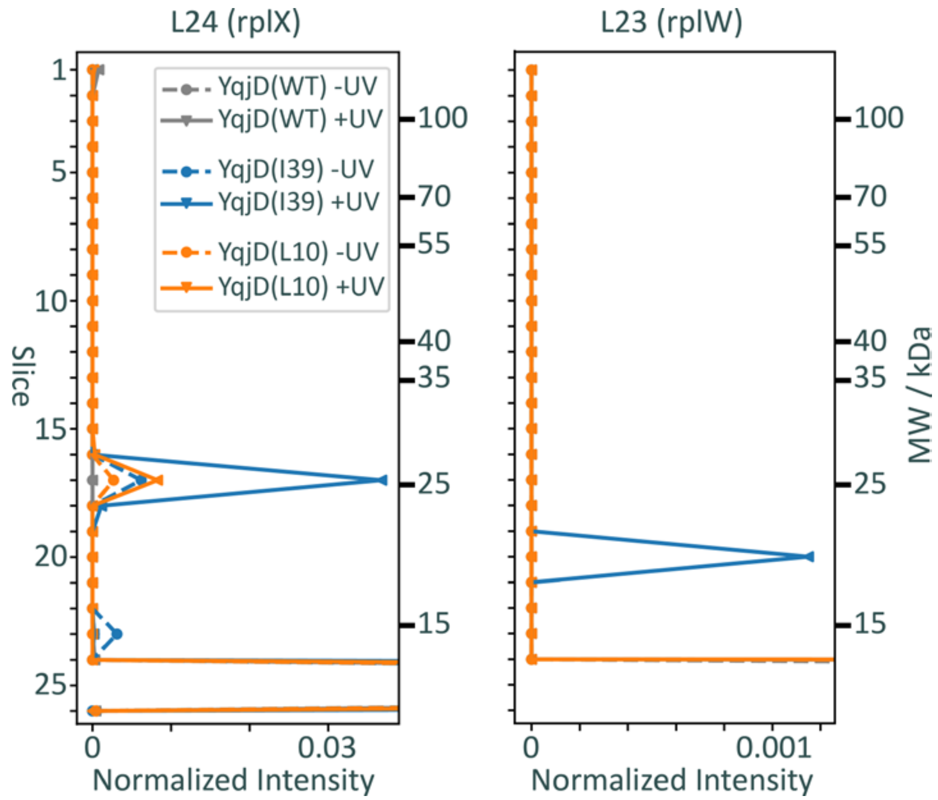


B

| Fraction | Ribosomes | | | | | | | | Crude membrane | | | | | | | |
|---|---|---|---|---|---|---|---|---|---|---|---|---|---|---|---|---|
| Ribosomes | G71pBpa | | | | E52pBPA | | | | G71pBpa | | | | E52pBPA | | | |
| Growth phase | Exp | | Stat | | Exp | | Stat | | Exp | | Stat | | Exp | | Stat | |
| UV | + | - | + | - | + | - | + | - | + | - | + | - | + | - | + | - |

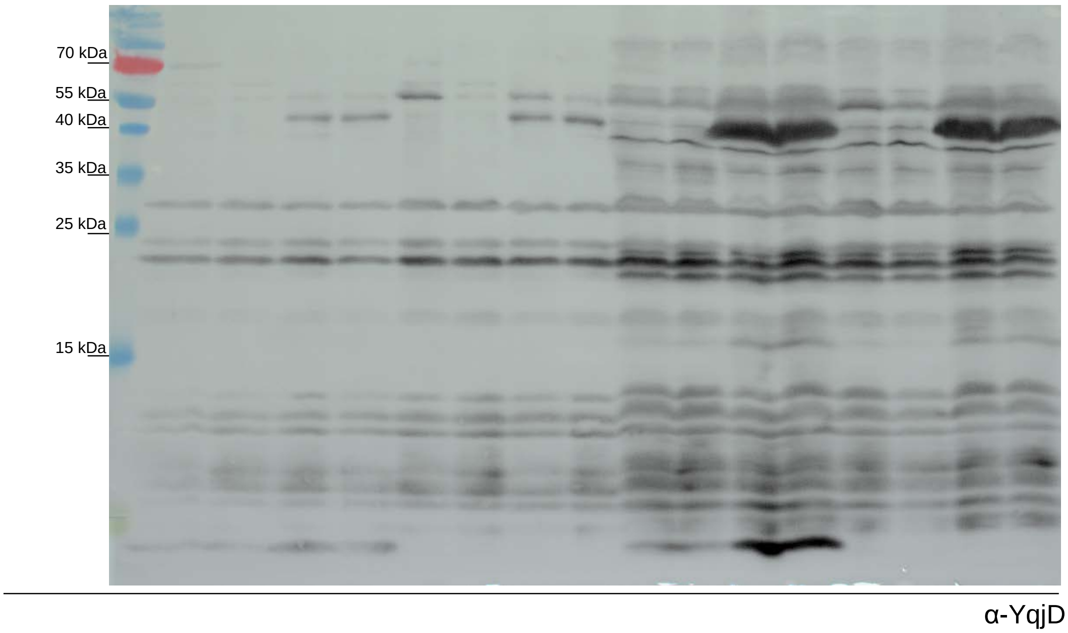


(Njenga et al., Supp. Fig. 7)
